# Foxp3 and BATF cooperatively direct *cis*-regulatory programs and gene expression for functional differentiation of Treg cells

**DOI:** 10.1101/2025.08.13.669552

**Authors:** Ryuichi Murakami, Norihito Hayatsu, Takahisa Miyao, Zhenan Liu, Shumpei Shibata, Masashi Matsuda, Yusuke Iizuka, Hideyuki Yoshida, Yoshinori Hasegawa, Wataru Ise, Haruhiko Koseki, Tomohiro Kurosaki, Shohei Hori

## Abstract

Mechanisms by which diverse transcription factors (TFs) shape the heterogeneous transcriptional and epigenetic landscape of regulatory T (Treg) cells remain poorly understood. By investigating interactions between BATF and Foxp3 TFs, we discovered their cooperative roles in directing *cis*-regulatory programs and gene expression essential for differentiation of immunosuppressive effector Treg (eTreg) cells. Simultaneous single-cell chromatin accessibility and transcriptome profiling, combined with topic modeling, identified *cis*-regulatory elements and associated programs jointly regulated by these TFs in eTreg cells. Genome-wide mapping of Treg cell-specific BATF and eTreg cell-specific Foxp3 binding sites revealed their co binding at some of these *cis*-elements, synergistically enhancing accessibility and transcription. Furthermore, we provide evidence that Foxp3 interacts with specific TFs to orchestrate diverse *cis*-regulatory programs among Treg cell differentiation states. Thus, Foxp3 serves as a master, but context-dependent regulator, cooperating with other TFs, including BATF, to shape the heterogeneous *cis* regulatory and transcriptional landscape critical for functional Treg cell differentiation.

## INTRODUCTION

Cellular differentiation and function are orchestrated by intricate interactions between transcription factors (TFs) and *cis*-regulatory elements (CREs), particularly enhancers. These CREs control gene expression by recruiting *trans* acting factors, including TFs, forming *cis*-regulatory programs that dictate cell identity and function. Among TFs, “master regulators” are pivotal, being necessary and sufficient for lineage specification^1^. However, how master regulators orchestrate these programs remains unclear. Do they opportunistically bind pre accessible CREs to modulate gene expression, or do they actively promote CRE accessibility in response to extracellular cues^2,3^?

Regulatory T (Treg) cells expressing the forkhead (FKH) TF, Foxp3, constitute a distinct lineage committed to immunosuppressive functions essential for immune tolerance^4^. Foxp3 is considered a master regulator, as Foxp3 mutations impair development of functional Treg cells, while ectopic expression converts Foxp3-CD4^+^ conventional T (Tconv) cells into a Treg cell-like phenotype^5–7^. However, T cells that transcribe the *Foxp3* locus, but fail to express functional protein, do develop, and exhibit a partial Treg cell-like phenotype, despite lacking suppressive function^8,9^. Conversely, Foxp3-transduced Tconv cells fail to fully acquire Treg cell transcriptional features^10,11^. These findings led to the conclusion that Foxp3-independent mechanisms, including epigenetic regulation and additional TFs, are required for Treg cell identity^12,13^.

Treg cells are heterogeneous in origin and function. Many are thymus-derived Treg cells, reportedly expressing Helios, while others are peripherally generated Treg (pTreg) cells, including Helios-RORγt^+^ pTreg cells induced by commensal bacteria^14^. Functionally, Treg cells are categorized as CD44^low^CD62L^high^CCR7^high^ central Treg (cTreg) or CD44^high^CD62L^low^CCR7^low^ effector Treg (eTreg) cells^15–17^. Like naïve Tconv (Tn) cells, cTreg cells circulate through secondary lymphoid tissues, where T cell receptor (TCR) activation drives their differentiation into eTreg cells that suppress immune responses in lymphoid and non-lymphoid tissues. eTreg cells also support tissue repair and maintenance^18^. Single-cell chromatin accessibility analyses have delineated these Treg cell states and implicated diverse TFs in shaping them^19^. However, mechanisms by which these TFs, particularly Foxp3, shape chromatin landscapes remain poorly understood.

Genome-wide analyses of chromatin accessibility and Foxp3 binding sites in Treg cells showed that Foxp3 predominantly occupies open chromatin regions (OCRs), pre-accessible in Tconv cells and established by other TFs^20^. Studies of Foxp3-deficient Treg-like cells further showed that Foxp3-dependent chromatin changes primarily occur at OCRs unbound by Foxp3^19,21^. These findings seemed to suggest that Foxp3 acts opportunistically, indirectly shaping the chromatin landscape by regulating intermediate TFs^20,21^. However, how Foxp3 regulates *cis* regulatory programs remains unclear, as prior studies did not identify functional OCRs and Foxp3 binding sites affecting gene expression. Additionally, Foxp3 binding sites were previously examined in total Treg cells without addressing heterogeneity^19–21^. Since many TF binding events are non-functional, and condition specific binding sites are enriched for functional events^22^, Foxp3 may bind and modulate CREs in a differentiation state-specific manner.

The AP-1 TF, BATF, is indispensable for differentiation of diverse effector lymphocytes, including effector Tconv (Teff) cell subsets, such as Th2, Th17, and Teh cells, and eTreg cells^23–28^. BATF heterodimerizes with JUN family proteins to bind DNA via AP-1 motifs and often partners with IRF4 to bind AP-1-IRF composite elements (AICEs)^23^. In T cells, BATF is upregulated by TCR signaling and is thought to act as a pioneer factor that opens chromatin and initiates epigenetic reprogramming^29–31^. While BATF functions in multiple lymphocyte populations, it promotes lineage^-^ or subset-specific gene expression, e.g., *Gata3* in Th2, *Il17* in Th17, and *Bcl6* in Teh cells^23,25^. However, whether and how it orchestrates Treg cell lineage-specific *cis*-regulatory programs remain unexplored.

In this study, by analyzing BATF-deficient Treg cells, we demonstrate that BATF promotes Treg cell-specific effector genes and OCRs. Examination of Treg-like cells expressing a DNA-binding-defective Foxp3 mutant revealed Foxp3’s critical role in enhancing BATF-dependent, eTreg cell-specific genes and OCRs. Simultaneous single-cell chromatin accessibility and transcriptome profiling identified putative CREs and *cis*-regulatory programs jointly regulated by BATF and Foxp3. Finally, mapping BATF and Foxp3 binding sites specific to Treg and eTreg cells and integrating these data with BATF^-^ and Foxp3-dependent CREs revealed that these TFs directly and cooperatively regulate *cis*-regulatory programs in eTreg cells.

## RESULTS

### Fatal multi-organ inflammation and eTreg cell deficiency result from BATF ablation in Treg cells

To investigate functions of BATF in Treg cells, we generated *Batf*^6l/6l^ *Foxp3*^YFPCre^ Treg cell-specific BATF conditional knock-out (cKO) mice. These mice spontaneously developed fatal multi-organ inflammation associated with a paucity of CD44^high^CCR7^low^ eTreg cells and accumulation of Teff cells, including Th1, Th2 and Th17 cells **(Figures S1A–S1E)**. To isolate cell-intrinsic effects of BATF deficiency in Treg cells in the absence of inflammation, we utilized *Batf*^6l/6l^ *Foxp3*^YFPCre/hCD2^ heterozygous female mice. Due to X-inactivation, these mice generate a mosaic of BATF-deficient YFP^+^hCD2^-^ cKO Treg cells and BATF-sufficient (wild type; WT) YFP^-^ hCD2^+^ Treg cells, of which the latter prevent disease development. BATF expression was specifically ablated in cKO Treg cells without affecting Foxp3 expression (**Figure S2A**). cKO Treg cells were deficient in CCR7^low^ eTreg, but not in CCR7^high^ cTreg cells in both lymphoid and non-lymphoid tissues **(Figure 1A)**. These findings underscore the critical role of BATF expression in Treg cells for maintaining self tolerance and for eTreg cell development.

**Figure 1:**
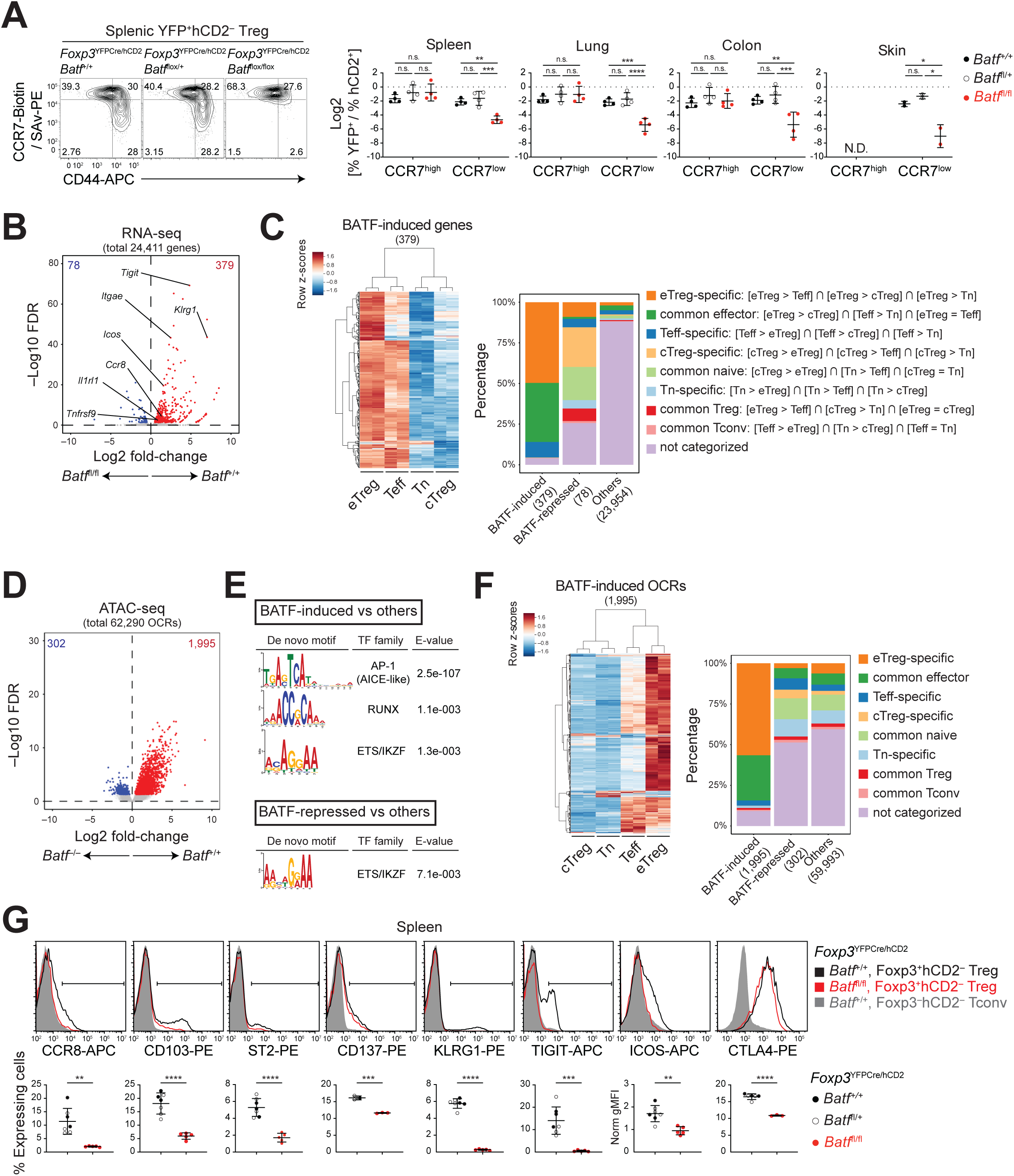
BATF promotes Treg cell-specific effector genes and OCRs. (A) FCM of BATF cKO and WT Treg cells. Graphs display log2 ratios of YFP^+^ to hCD2^+^ cells within CCR7^high^ or CCR7^low^ CD4^+^TCRβ^+^ populations (*n*=2–8; N.D., not detected in sufficient numbers). (B) RNA-seq analysis of cKO and WT Treg cells. Volcano plot comparing gene expression, highlighting BATF-induced (red) and BATF-repressed (blue) genes. (C) Heatmap illustrating expression levels of BATF-induced genes across CD4^+^ T cell subsets. Genes are categorized as indicated, with the percentage of each category summarized in stacked bar graphs. (D) ATAC-seq analysis of BATF KO and WT Treg cells. Volcano plot comparing chromatin accessibility, highlighting BATF-induced (red) and BATF-repressed (blue) OCRs. (E) Top three *de novo* motifs differentially enriched in each OCR category. (F) Heatmap showing accessibility of BATF-induced OCRs across CD4^+^ T cell subsets. Stacked bar graphs represent the percentage of each OCR category, defined as in (C). (G) FCM of cKO and WT Treg cells. Graphs show percentages of cells expressing indicated molecules or normalized geometric MFI (gMFI) of ICOS or CTLA4 in Foxp3^+^hCD2^-^ Treg cells (*n*=3–7). Data analyzed by two-way ANOVA with Sidak’s multiple comparisons test (A) or unpaired *t*-test (G). See also Figures S1 and S2.

### BATF promotes Treg cell-specific effector genes and OCRs

To determine whether BATF controls Treg cell-specific effector genes and/or common effector genes shared with Tconv cells, we performed RNA sequencing (RNA-seq) on cKO and *Batf*^−/–^ Treg (KO Treg) cells, as well as total Treg, cTreg, eTreg, Tn, and Teff cells from WT mice (**Figure S2B**). Differential expression analysis revealed genes either downregulated or upregulated in cKO Treg cells compared to WT Treg cells, designated as BATF-induced and -repressed genes, respectively **(Figure 1B, Table S1)**. Most BATF-induced genes were either specifically upregulated in eTreg cells (eTreg cell-specific genes) or shared with Teff cells (common effector genes) **(Figure 1C)**.

To investigate BATF effects on chromatin accessibility, we performed an assay for transposase-accessible chromatin with sequencing (ATAC-seq) of KO and WT Treg cells, as KO mice remain phenotypically healthy and their Treg cells are transcriptionally similar to those of cKO Treg cells **(Figures S2C, S2D)**. Differentially accessible OCRs were identified, with BATF-induced OCRs highly enriched for AICE-like AP-1 motifs **(Figures 1D, 1E, Table S2)**, suggesting direct regulation by BATF, partly in cooperation with IRF4. These BATF-induced OCRs primarily included eTreg cell-specific and common effector OCRs **(Figure 1F)**.

eTreg cell-specific and BATF-induced genes included *Ccr8*, *Icos*, *Ikzf2* (encoding Helios), *Il1rl1* (ST2), *Itgae* (CD103), *Klrg1*, *Tigit*, and *Tnfrsf9* (CD137) **(Table S1)**, reportedly upregulated in eTreg cells, including tissue-resident Treg cells, with some contributing to their differentiation and/or homeostasis^15–18,32,33^. *Ctla4 and Il10*, essential for Treg cell function^34,35^, were identified among genes associated with eTreg cell-specific and BATF-induced OCRs **(Table S2)**. Flow cytometry (FCM) analysis further validated BATF-dependent and eTreg cell-specific expression of selected molecules, including CTLA-4 **(Figures 1G, S2E)**.

Together, these results indicate that BATF can promote expression and chromatin accessibility of eTreg cell-specific genes and OCRs, thereby orchestrating lineage-specific programs critical for eTreg cell development and function.

### Foxp3 facilitates BATF-dependent eTreg cell-specific gene expression and chromatin accessibility

Foxp3-regulated or -bound genes were enriched in eTreg cell-specific BATF induced genes and OCRs (**Figure S2F**), leading us to hypothesize that Foxp3 is required for BATF to promote Treg cell lineage-specific effector programs. To test this hypothesis, we used *Foxp3*^R397W:hCD2^ mice, which carry the R397W mutation that fully eliminates regulatory functions of Foxp3 by disrupting its DNA-binding activity^26^ and examined whether mutant Treg-like cells resemble BATF-deficient Treg cells.

Treg-like cells from *Foxp3*^R397W:hCD2/GFP^ heterozygous mice (hereafter, R397W Treg cells) showed a selective eTreg cell deficiency in various tissues, although cTreg cells were also affected (**Figure 2A**). RNA-seq and ATAC-seq analysis of R397W Treg and WT Treg cells identified Foxp3-induced genes and OCRs (**Figure 2B, Table S3, S4**), most of which were eTreg cell-specific and common effector genes and OCRs (**Figure 2C**). Both expression and chromatin accessibility of most BATF-induced genes and OCRs were reduced in R397W Treg cells (**Figure 2D**), with eTreg cell-specific genes and OCRs particularly affected (**Figure 2E**), suggesting co regulation by BATF and Foxp3. Representative eTreg cell-specific BATF-induced molecules were indeed downregulated in R397W Treg cells (**Figure 2F**).

**Figure 2:**
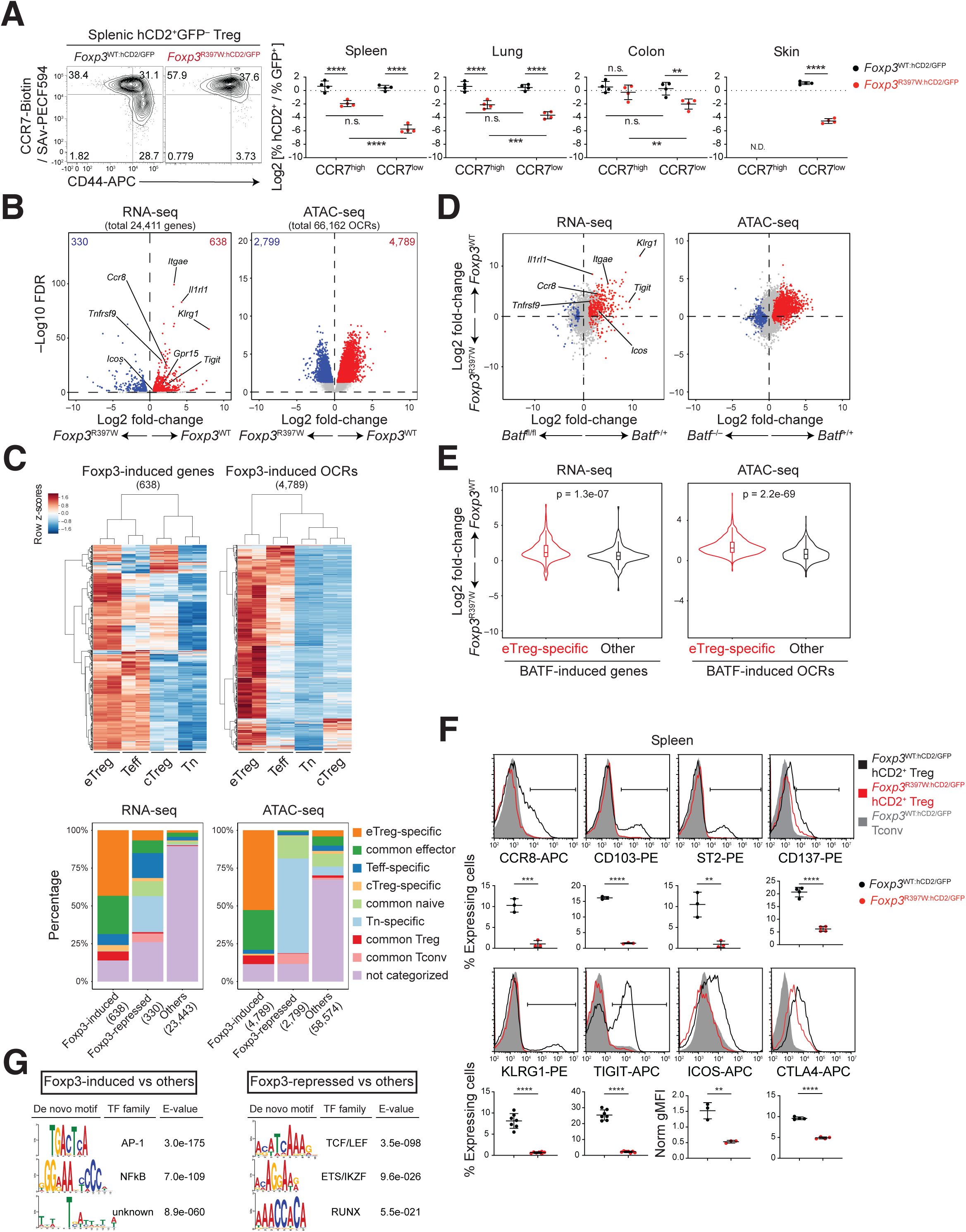
Phenotypic, transcriptional, and epigenetic commonalities between BATF-deficient and Foxp3 R397W Treg cells. (A) FCM of R397W and WT Treg cells. Graphs display log2 ratios of hCD2^+^ to GFP^+^ cells within CCR7^high^ or CCR7^low^ CD4^+^TCRβ^+^ populations (*n*=4). (B) RNA-seq and ATAC-seq analysis of R397W and WT Treg cells. Volcano plots comparing gene expression and chromatin accessibility, highlighting Foxp3-induced (red) and Foxp3-repressed (blue) genes and OCRs. (C) Heatmaps of expression levels and chromatin accessibility of Foxp3-induced genes and OCRs across CD4^+^ T cell subsets. Stacked bar graphs show the percentage of each gene or OCR category, defined as in Figures 1C and 1F. (D) FC vs. FC plots of RNA-seq and ATAC-seq data, comparing effects of BATF deficiency and the R397W mutation on gene expression and chromatin accessibility in Treg cells, highlighting BATF-induced (red) and BATF-repressed genes and OCRs (blue). (E) Log2 FC of gene expression and chromatin accessibility in WT vs. R397W Treg cells for indicated gene and OCR categories. (F) FCM of R397W and WT Treg cells. Graphs show percentages of cells expressing indicated molecules or normalized gMFI of ICOS or CTLA-4 in hCD2^+^ Treg cells (*n*=3– 7). Treg cells were gated as Foxp3^+^hCD2^+^ cells for intracellular CTLA-4 staining or as hCD2^+^GFP^-^ cells for other analyses. (G) Top three *de novo* motifs differentially enriched in each OCR category. Data analyzed by two-way ANOVA with Sidak’s multiple comparisons test (A), Mann-Whitney U test (E), or unpaired *t*-test (F). See also Figure S3.

We also generated a *Foxp3*^null^ (*Foxp3*^KO:hCD2^) allele with a frameshift mutation in exon 2 of the *Foxp3*^hCD2^ allele, which abolished Foxp3 protein expression and induced a *scurfy*-like disease in male mice (**Figures S3A-S3C**). Foxp3-deficient Treg like cells mirrored R397W Treg cells in exhibiting a selective eTreg cell deficiency, as well as comparable transcriptome and chromatin accessibility profiles (**Figures S3D, S3E**). Thus, effects observed in R397W Treg cells stem from loss of Foxp3 activity.

Differential motif enrichment analysis revealed AP-1 and NFκB motifs in Foxp3-induced OCRs, suggesting that Foxp3 relies on these TFs to promote chromatin accessibility (**Figure 2G**). Substantial overlap between BATF-induced and Foxp3-induced OCRs (**Figures 2D**, **2E**) highlights BATF as a critical AP-1 TF that Foxp3 utilizes to drive chromatin accessibility. Additionally, we identified many Foxp3-repressed OCRs, the majority of which were Tn cell-specific and enriched for TCF/LEF, ETS/IKZF, and RUNX motifs (**Figures 2B**, **2C**, **2G**).

Together, these findings demonstrate that Foxp3 promotes BATF-dependent, eTreg cell-specific gene expression and chromatin accessibility, suggesting functional cooperation between these TFs in regulating Treg cell effector programs.

### BATF and Foxp3 synergistically promote eTreg cell differentiation in response to TCR signaling

To assess whether BATF and Foxp3 cooperate to promote eTreg cell differentiation, we generated *Batf*^−/–^ *Foxp3*^R397W^ mice, isolated their Treg-like cells, and reconstituted them by retroviral transduction with Foxp3, BATF, both, or neither. This approach allowed us to evaluate whether these TFs could individually or jointly rescue eTreg cell differentiation. R397W Treg cells expressed lower levels of Foxp3 protein than WT Treg cells and lacked *in vitro* suppressive activity, both of which were fully restored by Foxp3 transduction (**Figures S4A, S4B)**. Thus, endogenous Foxp3 R397W protein does not interfere with the function of transduced WT Foxp3 protein.

These transduced cells were transferred into irradiated mice to assess their behavior *in vivo* (**Figures 3A**, **3B, S4C**). Foxp3 transduction alone had no impact on cell numbers in host tissues, proliferation, or expression of eTreg and Teff cell associated molecules. In contrast, BATF transduction alone induced proliferation and a common effector phenotype (characterized by CCR7 downregulation and CD44 upregulation), but failed to promote an eTreg cell phenotype, as defined by induction of CCR8, CD103, CTLA-4, ICOS, and ST2. Remarkably, co-transduction of BATF and Foxp3 resulted in robust increases in cell numbers, proliferation, and expression of these eTreg cell markers.

**Figure 3:**
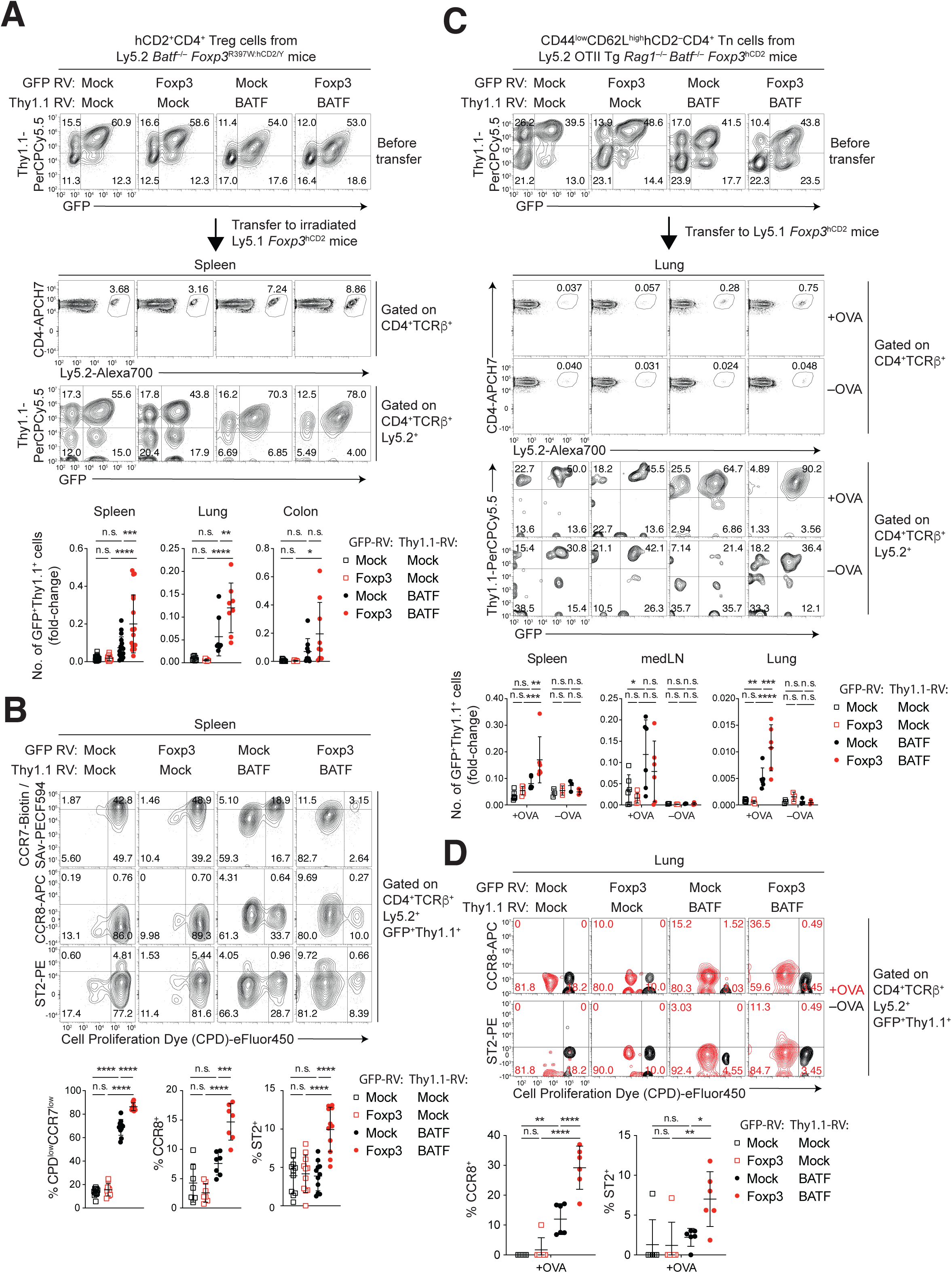
BATF and Foxp3 synergistically promote eTreg cell differentiation in response to TCR signals. (A, B) Ly5.2 *Batf*^−/–^ *Foxp3*^R397W^ Treg cells were transduced with the indicated GFP and Thy1.1 retroviruses (RV), labeled with Cell Proliferation Dye (CPD) eFluor 450, and transferred into irradiated Ly5.1 mice. On day 5 post-transfer, leukocytes were analyzed by FCM. (C, D) Ly5.2 OTII TCR transgenic *Batf*^−/–^ *Rag1*^−/–^ Tn cells were retrovirally transduced, CPD labeled, and transferred into non-irradiated Ly5.1 mice. On day 1, OVA or PBS was administered intranasally. On day 4, leukocytes were analyzed by FCM (medLN: mediastinal lymph node). (A, C) FCM of donor CD4^+^ T cells before and after transfer. Graphs show numbers of GFP^+^Thy1.1^+^ donor cells relative to transferred GFP^+^Thy1.1^+^ cells (*n*=8–18 for A, *n*=3–6 for C). (B, D) FCM of GFP^+^Thy1.1^+^ donor cells post-transfer. Graphs show percentages of cells with indicated phenotypes (*n*=7–11 for B, *n*=3–6 for D). Data analyzed by two-way ANOVA with Tukey’s multiple comparisons test (C) or one-way ANOVA with Tukey’s multiple comparisons test (A, B, D). See also Figure S4.

To determine whether TCR signaling is required for cooperative effects of BATF and Foxp3, we transduced BATF-deficient monoclonal Tn cells expressing the OTII TCR and transferred them into non-irradiated mice treated intranasally with or without the cognate antigen, ovalbumin (OVA). OTII T cells co-transduced with BATF and Foxp3 accumulated in greater numbers in the spleen and lung than those transduced with either TF alone or neither, but only in OVA-treated mice (**Figure 3C**). Additionally, BATF and Foxp3 co-transduction synergistically upregulated CCR8 and ST2 in OVA-treated mice (**Figure 3D**).

These findings demonstrate that BATF and Foxp3 act synergistically to drive eTreg cell differentiation, a process dependent on TCR signaling and characterized by induction of eTreg cell markers.

### BATF and Foxp3 shape the single-cell chromatin landscape of eTreg cells

To explore how BATF and Foxp3 may jointly regulate eTreg cell-specific gene expression and chromatin accessibility at the single-cell level, we performed paired scATAC-seq and scRNA-seq (scMultiome) analysis on WT Tconv, WT Treg, BATF KO Treg, and R397W Treg cells. Focusing on chromatin accessibility profiles to capture heterogeneity, we filtered and integrated these data with batch effect correction, performed dimensionality reduction and clustering, and excluded outliers **(Figures S5A-S5D)**. Re-clustering of 45,158 cells yielded 8 clusters, with no evidence of batch effect-driven clustering (**Figures 4A**, **4B**). WT Treg and Tconv cells were epigenetically distinct, segregating in the uniform manifold approximation and projection (UMAP) space. Clusters 0 and 7 were predominantly Tconv cells, whereas the remaining clusters were composed mainly of Treg cells of different genotypes, except cluster 4, which included both (**Figures 4B**, **4C**). BATF deficiency depleted cluster 3 and enriched cluster 6, whereas the R397W mutation depleted clusters 1 and 3 while enriching cluster 2.

**Figure 4:**
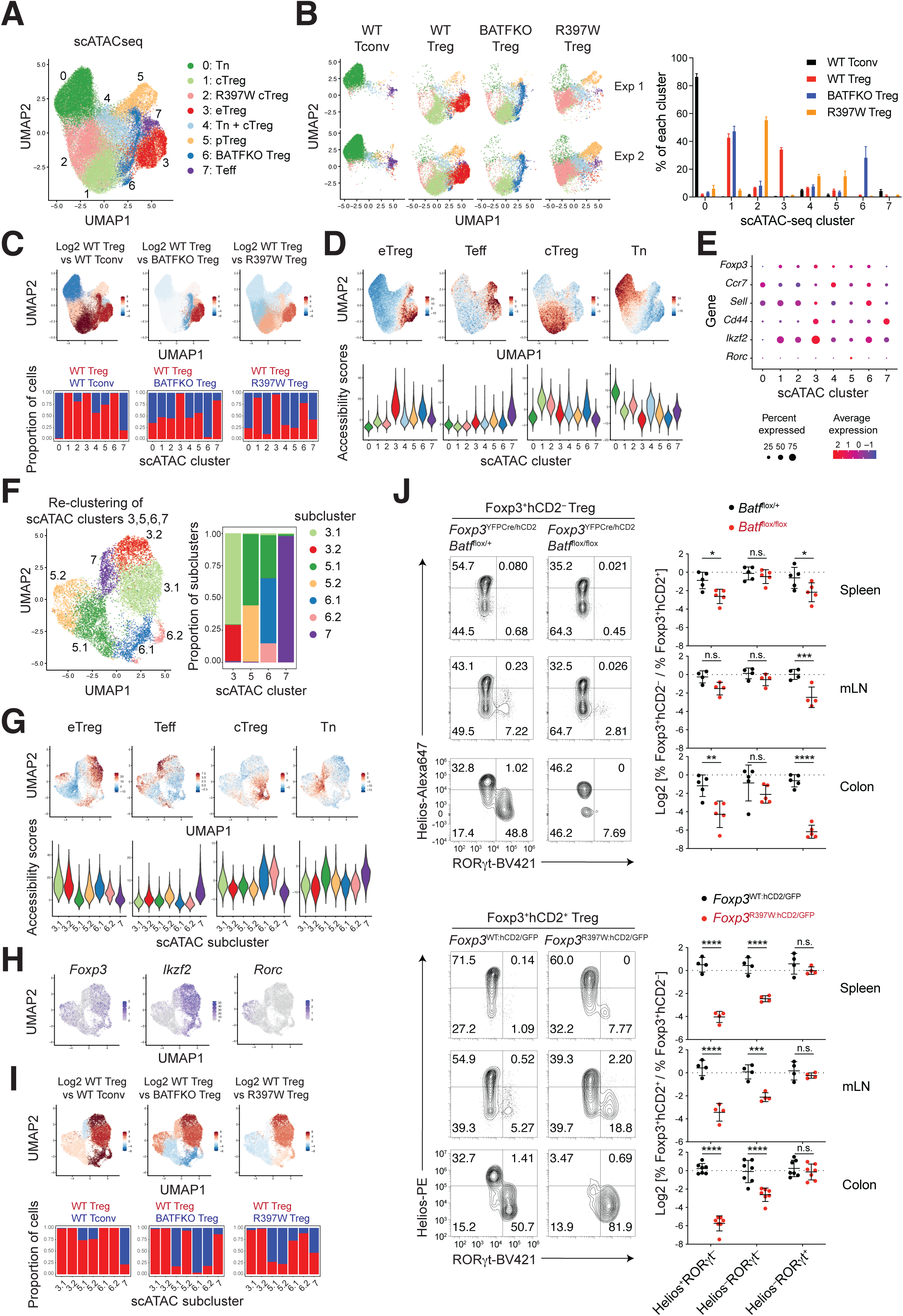
scMultiome analysis of WT Tconv, WT Treg, BATF KO Treg, and R397W Treg cells. (A) Merged UMAP of integrated scATAC-seq data. (B) UMAP showing individual samples from two independent experiments. Graph summarizes cluster percentages. (C) Clusters color-coded by the log2 ratio of their average frequency in WT Treg cells vs. WT Tconv, BATF KO Treg, or R397W Treg cells, visualized on UMAP. Graphs show average proportions of WT Treg cells (red) and other populations (blue) per cluster. (D) Accessibility scores for CD4^+^ T cell subset-specific OCRs in single cells (UMAPs) or their distribution per cluster (violin plots). (E) Cluster-level expression of indicated genes. (F-I) Re-clustering and analysis of eTreg and Teff cell clusters 3 and 5–7. (F) Merged UMAP of re-clustered scATAC-seq data. Stacked bar graphs show subcluster proportions per cluster. (G) Accessibility scores for CD4^+^ T cell subset-specific OCRs. (H) Single-cell expression of indicated genes. (I) Subclusters color-coded and visualized, with graphs summarized as in (C). (J) FCM of BATF cKO and R397W Treg cells and their WT controls. Graphs display log2 ratios of Foxp3^+^hCD2^-^ to Foxp3^+^hCD2^+^ cells (BATF cKO) or Foxp3^+^hCD2^+^ to Foxp3^+^hCD2^-^ cells (R397W) in indicated subsets (*n*=4–7). Data analyzed by two-way ANOVA with Tukey’s multiple comparisons test (J). See also Figure S5.

To annotate these clusters, we quantified marker gene expression and accessibility of CD4^+^ T cell subset-specific OCRs defined by bulk ATAC-seq (**Figures 4D**, **4E, S5E**). Clusters 0, 1, 3, and 7 corresponded to Tn, cTreg, eTreg, and Teff cells, respectively, based on high accessibility of subset-specific OCRs. Cluster 2, enriched in R397W Treg cells, was designated as R397W cTreg cells, due to its high cTreg cell specific OCR accessibility. Cluster 4 was a mixture of Tn and cTreg cells, albeit more activated with higher *Cd44* expression. Cluster 5 represented pTreg cells with low *Ikzf2* (Helios) expression, some expressing *Rorc* (RORγt) and gene signatures of pTreg cells^36,37^ (**Figures 4E, S5E, S5F**). Cluster 6, primarily found in BATF KO Treg cells, was annotated accordingly.

Beyond cluster 3, fractions of clusters 5 and 6 also exhibited high eTreg cell specific chromatin accessibility (**Figure 4D**). To refine the definition of eTreg cell subpopulations, we re-clustered clusters 3 and 5-7, identifying 7 subclusters (**Figure 4F**). Since these subclusters were included in the parent clusters, except for one that spans clusters 5 and 6 (designated 5.1), we referred to subclusters included in cluster *x* as subclusters *x*.1, *x*.2, and so on. Clusters 3 (both 3.1 and 3.2), 5.2, and 6.1, corresponded to eTreg cells (**Figure 4G**). However, a fraction of cluster 6.1 showed higher cTreg cell-specific OCR accessibility, suggesting a less differentiated state. *Ikzf2*, *Rorc*, and pTreg cell gene signatures identified cluster 5.2 as peripherally generated RORγt^+^ eTreg cells, whereas clusters 3 and 6.1 corresponded to thymus derived Helios^+^ eTreg cells (**Figures 4E**, **4H, S5G**). Cluster 6.1 was composed exclusively of BATF KO Treg cells, and cluster 5.2 was absent in BATF KO, but not in R397W Treg cells (**Figure 4I**). Indeed, RORγt^+^ Treg cells were deficient in BATF cKO, but not in R397W Treg cells, whereas Helios^+^ Treg cells were depleted in both BATF cKO and R397W Treg cells, consistent with cluster 3 deficiency (**Figure 4J**). However, Helios^+^ Treg cells were less affected in BATF KO Treg cells, likely due to expansion of cluster 6.

These findings highlight distinct and overlapping roles of BATF and Foxp3 in shaping the chromatin landscape of Treg cells. Both TFs are essential for generation of Helios^+^ eTreg cells, whereas BATF alone is required for RORγt^+^ eTreg cells. Additionally, BATF deficiency produces a unique Helios^+^ eTreg cell population absent in WT Treg cells, and Foxp3 globally regulates chromatin accessibility in cTreg cells.

### Chromatin features regulated by BATF and Foxp3

To characterize OCRs regulated by BATF and Foxp3, we compared aggregated scATAC-seq data from WT Treg, BATF KO Treg, and R397W Treg cells and identified OCRs induced or repressed by these TFs. BATF^-^ and Foxp3-induced OCRs showed substantial overlap (**Figure 5A**). High accessibility of these overlapping OCRs in Helios^+^ eTreg cell cluster 3 (3.1 and 3.2) and RORγt^+^ eTreg cell cluster 5.2 indicates that BATF and Foxp3 jointly enhance chromatin accessibility in these eTreg cell subsets (**Figure 5B**, **5C**). BATF also promoted other OCRs selectively accessible in these clusters and Teff cluster 7, independently of Foxp3.

**Figure 5:**
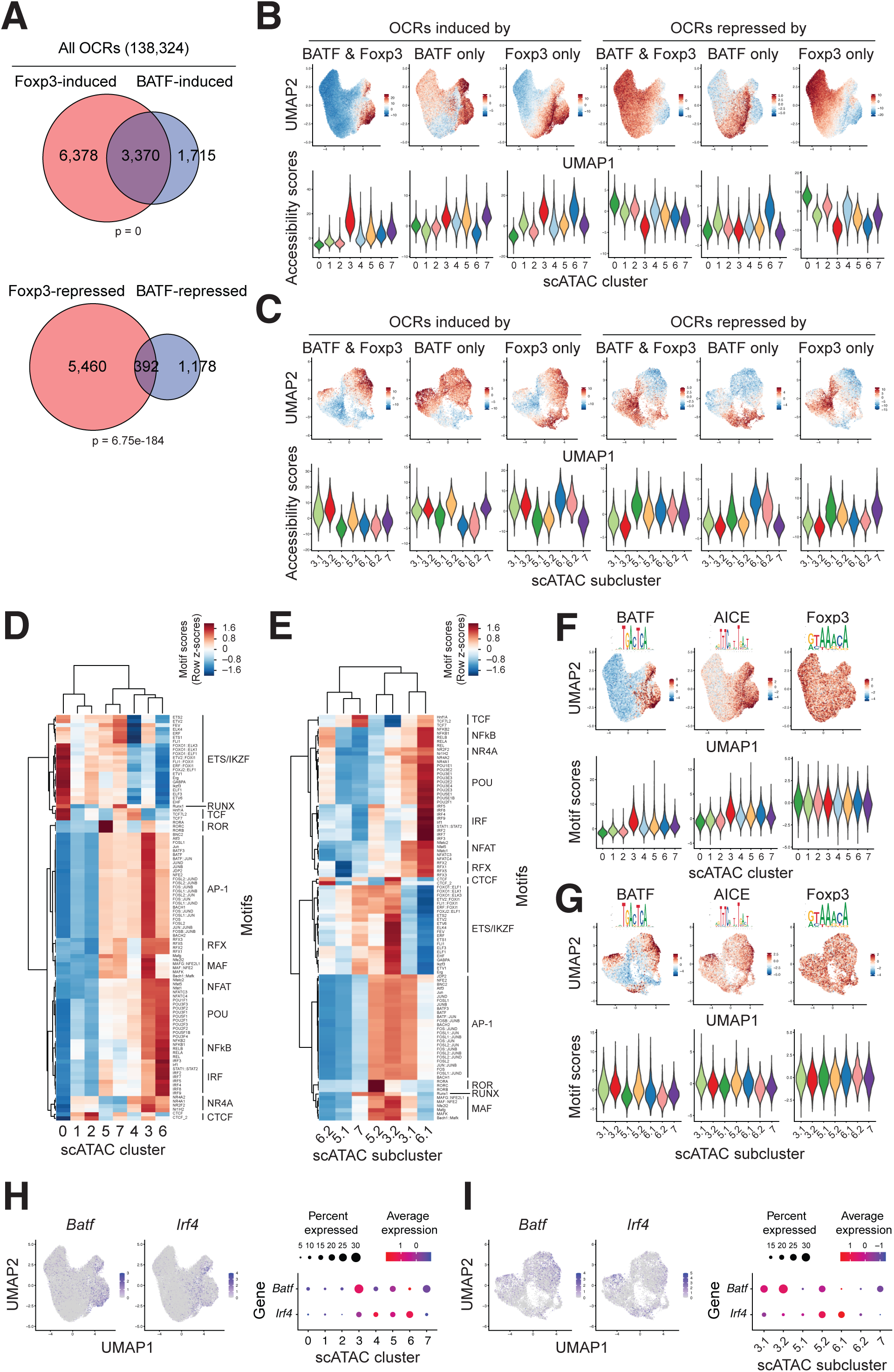
Single-cell characteristics of OCRs regulated by BATF and Foxp3. (A) Venn diagrams showing overlap between OCR groups. *P*-values are from hypergeometric tests. (B, C) Accessibility scores for indicated OCR groups in single cells for all cells (B) or eTreg and Teff cell subclusters (C) (UMAPs), and their distribution per cluster (B) or subcluster (C) (violin plots). (D, E) Heatmaps showing TF motif activity scores for the 100 most variable motifs (defined in Figure S5H), averaged per cluster (D) or subcluster (E). (F, G) TF motif activity scores for indicated motifs in single cells for all cells (F) or eTreg and Teff cell subclusters (G) (UMAPs), and their distribution per cluster (F) or subcluster (G) (violin plots). (H, I) Expression of indicated genes at the single-cell or cluster level for all cells (H) or eTreg and Teff cell subclusters (I). See also Figure S5.

We quantified TF motif activities using chromVAR^38^. BATF and other AP-1 motif activity scores were highest in clusters 3 and 5.2, suggesting that BATF directly increases chromatin accessibility at BATF-induced OCRs (**Figures 5D-5G**). Interestingly, BATF scores were lower in cluster 7 (**Figure 5G**), despite high accessibility of BATF-induced OCR (**Figure 5C**) and substantial *Batf* expression (**Figures 5H**, **5I**), implying that Foxp3 enhances BATF activity in clusters 3 and 5.2. Additionally, AICE motif scores and *Irf4* expression were also detected across eTreg cells, albeit more broadly than BATF scores and *Batf* expression (**Figures 5F-5I**). In BATF KO eTreg cell cluster 6.1, motif activities of non-AP-1 TFs associated with T cell activation, such as NFκB, NFAT, and NR4A, were elevated (**Figure 5E**). This suggests that in the absence of BATF, Helios^+^ eTreg cells are partially generated by actions of these TFs, but remain in a less differentiated state (cluster 6.1) without progressing to the fully differentiated state of cluster 3.

Foxp3 facilitated accessibility in clusters 3, 6.1, and 6.2 independently of BATF (**Figures 5B**, **5C**). Notably, Foxp3 motif activities were uniform across single cells, suggesting that Foxp3 alone is insufficient to drive chromatin accessibility (**Figures 5F, 5G, S5H**). Instead, Foxp3 appears to rely on NFκB, NFAT, and NR4A to promote chromatin accessibility in cluster 6.1 and collaborates with BATF to achieve full differentiation in cluster 3 (**Figures 5D**, **5E**).

In contrast, Tn cell cluster 0 exhibited the highest accessibility of Foxp3-repressed OCRs and elevated motif activities of TCF/LEF, ETS/IKZF, and RUNX TFs (**Figures 5B**, **5D**). Similarly, Teff cell cluster 7 displayed greater accessibility of these OCRs and higher motif activities for these TFs than eTreg cell clusters (**Figures 5C**, **5E**). Taken together with bulk ATAC-seq results (**Figures 2C**, **2G**), these findings suggest that Foxp3 represses OCRs preferentially accessible in Tn and Teff cells through interactions with these TFs. Notably, OCR accessibility and motif activities in R397W cTreg cell cluster 2 were intermediate between those of cluster 0 and cTreg cell cluster 1 (**Figures 5B**, **5D**), reelecting both Foxp3-dependent and ^-^ independent repression mechanisms.

Taken together, these findings reveal a dual role of Foxp3 in Treg cells, repressing chromatin features characteristic of Tconv cells while promoting eTreg cell-specific OCR accessibility. Foxp3 appears to dynamically interact with specific TFs, including BATF, to shape chromatin states according to their differentiation state, driving Treg cell lineage-specific chromatin accessibility.

### Cis-regulatory programs regulated by BATF and Foxp3

To identify *cis*-regulatory programs regulated by BATF and Foxp3, we analyzed scMultiome data to identify putative *cis*-regulatory elements (CREs) among the 138,324 OCRs detected, using an established analytical framework^39,40^. By correlating chromatin accessibility with gene expression in single cells, we identified 7,608 OCR-gene links, comprising 6,240 OCRs associated with 2,037 genes. Most links (99.6%) showed positive correlations, indicating enhancer or promoter activity. Linked OCRs were primarily enriched near transcription start sites (TSSs), but also included distal elements (**Figure S6A**). While most OCRs were linked to single genes, some genes were associated with multiple OCRs (**Figures 6A, S6B**). A prior study showed that lineage-defining genes often contain multiple OCRs overlapping with super-enhancers (SEs)^39^. Consistently, Treg or Tconv cell SE genes^41^ had more linked OCRs than other genes, as observed at the *Ctla4* and *Ikzf2* loci (**Figures 6A, S6C**). Additionally, enhancer-promoter loops identified in Treg and/or Tconv cells^42^ were significantly enriched among linked OCRs (**Figure S6D**). These findings validate OCR-gene links as predictors of CREs. Furthermore, OCRs regulated by BATF and/or Foxp3 were enriched among linked OCRs (**Figure 6B**), underscoring their roles in CRE accessibility.

**Figure 6:**
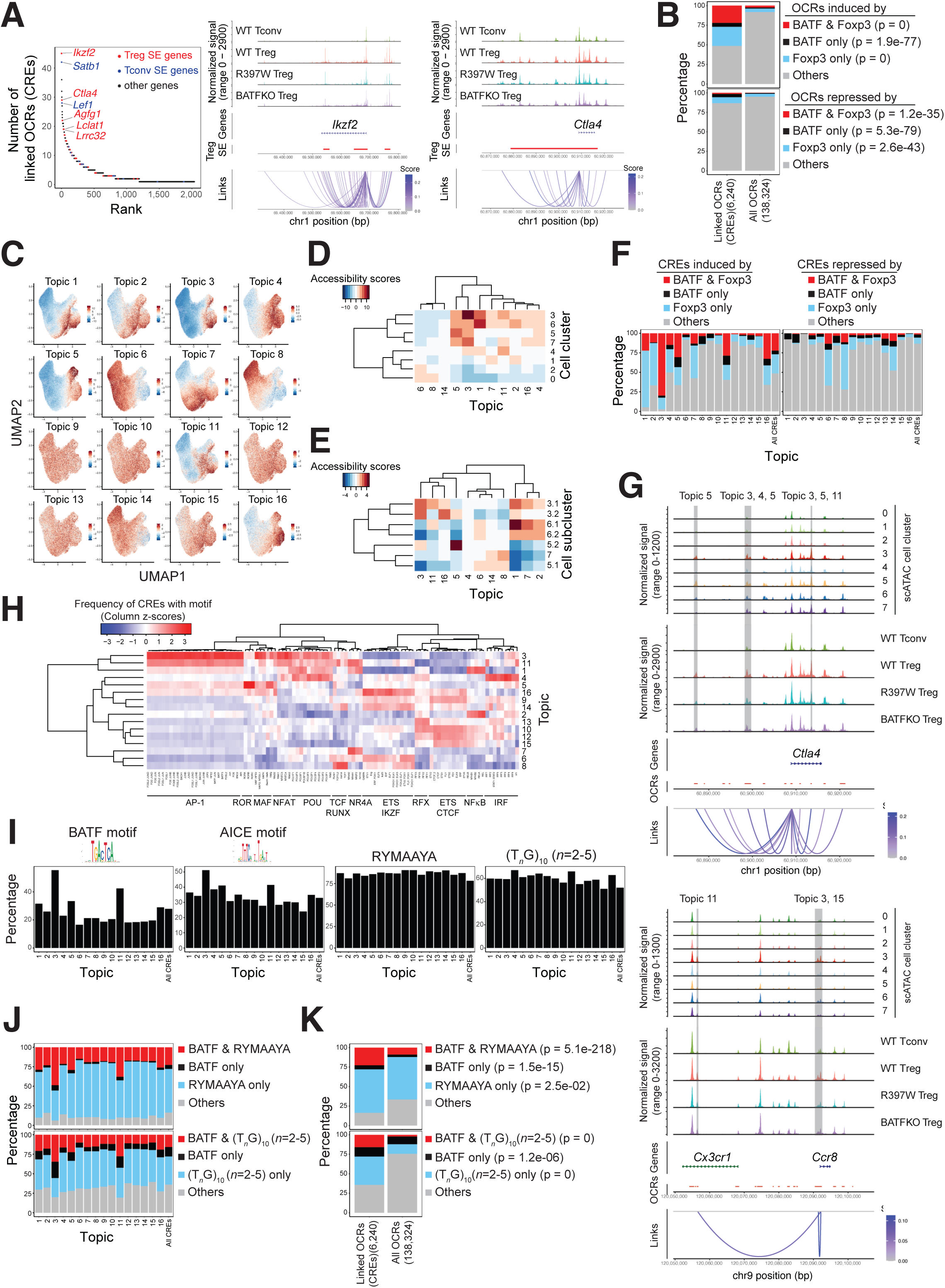
Cis-regulatory programs regulated by BATF and Foxp3. (A) Numbers of OCR-gene links per gene, ranked by abundance. Treg and Tconv SE genes^41^ are highlighted. Aggregated scATAC-seq tracks for indicated populations are shown at specific loci, with Treg SEs (red lines) and OCR-gene links (arcs). (B) Percentages of indicated OCR groups (defined in Figure 5A) among linked OCRs (CREs) or all OCRs. (C) Accessibility scores for topic-associated CREs, visualized on UMAPs for all cells. (D, E) Heatmaps of accessibility scores for CREs associated with variable topics, averaged per cluster (D) or subcluster (E). (F) Percentages of indicated CRE groups in topic-associated or all CREs. (G) Aggregated scATAC-seq tracks for indicated clusters or populations at specific loci, with OCR positions (red lines) and OCR-gene links (arcs). BATF^-^ and Foxp3-induced CREs associated with specific topics are highlighted in gray. (H) Heatmap of normalized frequency of CREs containing indicated TF motifs in topic-associated CREs. (I, J) Percentages of CREs with indicated (combinations of) TF motifs among topic associated or all CREs. (K) Percentages of OCRs with indicated combinations of TF motifs in CREs or all OCRs. Data analyzed by Fisher’s exact test (B, K). See also Figure S6.

To infer *cis*-regulatory programs as combinations of co-accessible CREs across single cells, we applied topic modeling to scATAC-seq data of linked OCRs (hereafter referred to as CREs) using cisTopic^43^. The optimal model yielded 16 topics, each associated with representative CRE sets (**Table S5**). While some topics showed minimal variability, others exhibited clear accessibility gradients in UMAP space (**Figures 6C, S6E**).

Based on hierarchical clustering of accessibility per cluster, variable topics can be categorized into those preferentially accessible in Tn and cTreg cell clusters 0-2 (topics 6, 8, and 14; category 1) and those preferentially accessible in eTreg and Teff cell clusters (**Figures 6C**, **6D**). The latter can be further categorized into two groups (**Figure 6E**). Topics 3, 5, 11, and 16 (category 2) represented eTreg cell programs, with topics 3, 11, and 16 enriched in Helios^+^ eTreg cell clusters 3.1 and/or 3.2, and topic 5-enriched in RORγt^+^ eTreg cell cluster 5.2 (**Figures 6E, S6E**). Topics 1, 2, and 7 (category 3) represented cTreg/“early” eTreg cell programs, enriched in BATF KO Treg cell clusters 6.1 and 6.2 (“early” Helios^+^ eTreg cells) alongside eTreg cell cluster 3.1, with topics 2 and 7 also enriched in cTreg cell cluster 1 (**Figures 6C-6E, S6E**). Notably, Teff cell cluster 7 showed enrichment for topics 8 and 14 compared to eTreg cell clusters (**Figures 6E, S6E**), suggesting shared *cis*-regulatory programs between Tn and Teff cells. Therefore, category 1 topics represent Tconv/cTreg cell programs.

Analysis of BATF deficiency and the R397W mutation revealed enrichment of BATF^-^ and Foxp3-induced CREs in topics of category 2, particularly topic 3 (**Figures 6F, S6F**), identifying eTreg cell programs co-regulated by these TFs. Such topics included those linked to eTreg cell-specific genes, such as *Ccr8*, *Ctla4*, *Icos*, *Ikzf2*, *Tigit*, and *Tnfrsf9* (**Figure 6G**, **Table S5**). BATF dependence was further supported by enrichment of AICE, BATF, and other AP-1 motifs in these topics, particularly topic 3 (**Figures 6H**, **6I**). Additional motif enrichments included MAF (topics 3 and 5), ROR (topic 5), NFAT (topics 3 and 11), NR4A (topic 11), and ETS/IKZF (topic 16) motifs, suggesting these combinations regulate distinct eTreg cell programs.

Foxp3 promoted category 3 programs largely independently of BATF (**Figures 6F, S6F**). Topics 1 and 2 were enriched for NFκB motifs, while topics 1 and 7 contained NR4A motifs (**Figure 6H**), suggesting that Foxp3 leverages these TFs to regulate cTreg/“early” eTreg cell programs. Conversely, category 1 programs (topics 6, 8, and 14) were enriched for Foxp3-repressed CREs and those containing TCF/LEF, ETS/IKZF, and RUNX motifs (**Figures 6F**, **6H**), implying that these TFs drive Tconv/cTreg cell programs that are counter-regulated by Foxp3 in Treg cells.

What about Foxp3 motifs? The canonical Foxp3 motif, determined by high throughput SELEX^44^, was detected only in a small fraction of CREs across all topics (**Figure S6G**). However, Foxp3 can bind a broader range of FKH motifs (RYMAAYA) and variant sequences through homo^-^ or hetero-dimerization^26,45–47^ and T*_n_*G repeats via homo-multimerization^48^. While not enriched in particular topics, these sequences appeared in most CREs (**Figure 6I**), with many CREs containing BATF or AICE motifs also containing FKH motifs or T*_n_*G repeats (**Figures 6J, S6H**). OCRs with both BATF or AICE motifs and FKH motifs or T*_n_*G repeats were enriched among CREs relative to all OCRs, suggesting a regulatory role for their co-occurrence (**Figures 6K, S6I**).

### BATF and Foxp3 directly and cooperatively promote eTreg cell cis-regulatory programs

To determine whether BATF and Foxp3 directly bind and promote accessibility of BATF^-^ and Foxp3-dependent CREs, we performed chromatin immunoprecipitation sequencing (ChIP-seq) for these TFs. Mapping BATF binding sites in Treg vs. Tconv cells and Foxp3 binding sites in eTreg vs. cTreg cells revealed both unique and common binding sites for each TF (**Figure 7A**). AP-1 motifs were enriched at BATF binding sites, particularly those overlapping CREs, whereas T*_n_*G repeats, FKH motifs, and motifs of Foxp3 partners, e.g., ETS/IKZF, RUNX, were enriched at Foxp3 binding sites (**Figures S7A, S7B)**, validating the ChIP-seq results.

**Figure 7:**
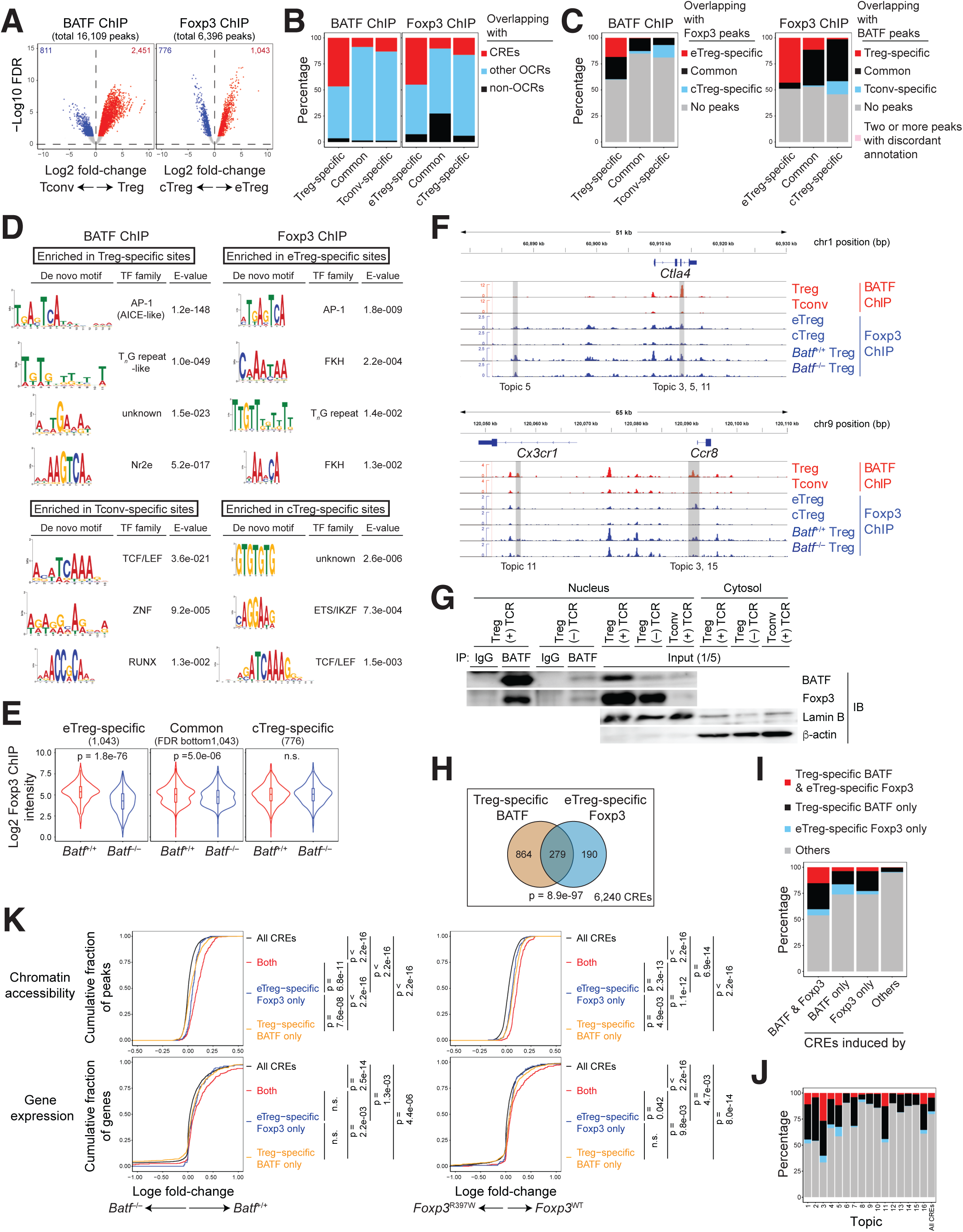
BATF and Foxp3 co-binding synergistically increases chromatin accessibility and gene expression. (A) Volcano plots comparing normalized read concentrations of BATF binding sites between Treg and Tconv cells, or Foxp3 binding sites between eTreg and cTreg cells. Differential binding sites are highlighted (red or blue). (B) Percentages of BATF or Foxp3 binding sites overlapping with CREs, other OCRs, or non-OCRs. (C) Percentages of BATF or Foxp3 binding sites overlapping with indicated categories of Foxp3 or BATF binding sites, respectively. (D) Top four *de novo* motifs differentially enriched at indicated binding sites. (E) Log2 normalized read concentrations of Foxp3 binding sites, comparing WT and BATF KO Treg cells in indicated categories. (F) BATF and Foxp3 ChIP-seq tracks. BATF^-^ and Foxp3-induced CREs overlapping with Treg-specific BATF and eTreg-specific Foxp3 binding sites are highlighted in gray. (G) Nuclear proteins from TCR-stimulated or unstimulated Treg cells were immunoprecipitated (IP) with anti-BATF or isotype-matched mAbs, followed by immunoblotting (IB). Representative results from two independent experiments. (H) Venn diagram showing overlap between Treg-specific BATF binding sites and eTreg-specific Foxp3 binding sites among CREs. (I, J) Percentages of CREs overlapping with specified categories of BATF and/or Foxp3 binding sites in indicated CRE categories (I) or topic-associated or all CREs (J). (K) Cumulative distribution function (CDF) plots showing effects of BATF deficiency or the R397W mutation on accessibility of indicated CRE categories and expression of associated genes. Data analyzed by Mann-Whitney U test (E), hypergeometric test (H), or Kolmogorov-Smirnov test (K). See also Figure S7.

Treg cell-specific BATF and eTreg cell-specific Foxp3 binding sites overlapped CREs more frequently than did common or Tconv/cTreg cell-specific sites (**Figure 7B**). BATF deficiency or the R397W mutation selectively impaired chromatin accessibility of CREs overlapping Treg cell-specific BATF or eTreg cell-specific Foxp3 binding sites, respectively, accompanied by reduced expression of linked genes (**Figure S7C**). These unshared binding sites are therefore enriched for functional regulatory events^22^.

Notably, Treg cell-specific BATF binding sites overlapped more frequently with eTreg cell-specific Foxp3 binding sites than did other BATF binding sites, and vice versa (**Figure 7C**). Enrichment of AICE-like AP-1 motifs and T*_n_*G repeats in Treg cell specific BATF binding sites, along with AP-1 and FKH motifs and T*_n_*G repeats in eTreg cell-specific Foxp3 binding sites, suggests their cooperative binding (**Figure 7D**). Foxp3 binding to eTreg cell-specific sites was impaired in BATF-deficient Treg cells, suggesting BATF-dependent recruitment of Foxp3 (**Figures 7E**, **7F**). Co immunoprecipitation assays confirmed physical interactions of BATF with Foxp3 in Treg cells, particularly upon TCR stimulation, likely reelecting their upregulation (**Figure 7G**). These results indicate that BATF and Foxp3 are recruited to the same genomic regions by binding to their recognition sequences and forming a molecular complex.

To examine functional effects of co-binding, we analyzed CREs with Treg cell specific BATF and eTreg cell-specific Foxp3 binding. These shared binding sites, enriched among CREs (**Figure 7H**), overlapped significantly with CREs containing both AP-1 or AICE motifs and FKH motifs or T*_n_*G repeats (**Figure S7D**), CREs induced by both BATF and Foxp3 (**Figure 7I**), and CREs associated with eTreg cell programs (topics 3, 5, 11, and 16) (**Figures 7J**), including loci such as *Ccr8*, *Ctla4*, *Icos*, and *Ikzf2* (**Figure 7F, Table S5**). Importantly, chromatin accessibility was more severely impaired by BATF deficiency or the R397W mutation for co-bound CREs compared to CREs bound by either TF alone, and this impairment was accompanied by reduced expression of linked genes (**Figure 7K**). This suggests that BATF and Foxp3 co binding synergistically enhances chromatin accessibility and gene expression. Together, these findings suggest that BATF and Foxp3 form a molecular complex that co-binds specific CREs in eTreg cells, directly and cooperatively enhancing chromatin accessibility and gene expression.

### cTreg cell-specific Foxp3 binding represses Tconv/cTreg cell cis-regulatory programs

We observed that cTreg cell-specific Foxp3 binding sites were enriched for ETS/IKZF and TCF/LEF motifs (**Figure 7D**). Chromatin accessibility of CREs overlapping these sites and expression of their linked genes were increased by the R397W mutation (**Figure S7C**). Furthermore, CREs associated with Tconv/cTreg cell programs (topics 6, 8, 14) were also enriched for cTreg cell-specific Foxp3 binding sites (**Figure S7E**). These results suggest that in cTreg cells, Foxp3 interacts with these TFs to occupy CREs that are preferentially accessible in Tconv and cTreg cells, repressing their chromatin accessibility and gene expression.

## DISCUSSION

Foxp3 serves as a master regulator of Treg cells, yet mechanisms underlying its regulatory actions remain incompletely understood. Here, we revisited the paradigm of Foxp3 as an opportunistic TF that indirectly influences the chromatin landscape^20,21^. By dissecting the interplay between Foxp3 and BATF at both population and single-cell levels, we demonstrate that they cooperatively and directly drive *cis*-regulatory programs and gene expression critical for Treg cell function. Our findings further suggest that Foxp3 changes its TF partners to orchestrate distinct *cis*-regulatory programs, highlighting its dynamic and versatile roles across Treg cell differentiation states.

We demonstrate that BATF-dependent development of eTreg cells is essential for maintaining self-tolerance. Previous studies reported increased Th2 cytokine production, altered triglyceride metabolism, and unstable Foxp3 expression in BATF cKO Treg cells^49,50^. However, because those analyses were conducted in diseased *Foxp3*^YFPCre^ *Batf*^6l/6l^ mice, these phenotypes likely reelect secondary consequences of inflammation rather than Treg cell-intrinsic defects. By analyzing non-inflammatory *Foxp3*^YFPCre/hCD2^ *Batf*^6l/6l^ mice, we unequivocally demonstrate the cell-intrinsic, non-redundant role of BATF in development of functional eTreg cells in both lymphoid and non-lymphoid tissues, supporting previous studies^26–28,51^.

BATF, expressed in multiple lymphocyte populations, upregulates lineage^-^ or subset-specific target genes^23^ presumably by facilitating chromatin accessibility at enhancers^29–31^. Consistently, we found that BATF promotes Treg cell-specific effector genes and OCRs. In Th17 and Tr1 cells, BATF collaborates with cytokine induced TFs, i.e., STAT3 for Th17 and IRF1 for Tr1, rather than master regulators, i.e, RORγt for Th17, to shape the epigenetic landscape^29,30^. In contrast, our findings identify Foxp3, the lineage-specific master regulator, as an essential BATF partner in Treg cells. Although Foxp3 was suggested to enhance eTreg cell-associated molecules by increasing BATF expression^51^, we find this unlikely, as BATF overexpression alone was insufficient. Foxp3 transduction was also required to induce an eTreg cell-like phenotype in BATF-deficient R397W Treg cells.

We demonstrated that both BATF and Foxp3 are required for development of Helios^+^ eTreg cells, whereas only BATF is required for RORγt^+^ eTreg cells. A previous study similarly reported the requirement of Foxp3 for Helios^+^ eTreg cells, but not RORγt^+^ eTreg cells; however, underlying mechanisms remain unexplored^19,37^. Our findings identify BATF as an essential partner of Foxp3 in shaping the chromatin landscape of Helios^+^ eTreg cells. Although not required for RORγt^+^ eTreg cells, Foxp3 facilitates accessibility of many BATF-dependent CREs shared between RORγt^+^ and Helios^+^ eTreg cells and upregulates associated genes. Thus, Foxp3 collaborates with BATF to promote *cis*-regulatory programs that operate in both eTreg cell subsets. Our results challenge the paradigm of Foxp3 as an opportunistic TF that indirectly influences the chromatin landscape of Treg cells^20,21^. Although previous studies suggested that Foxp3-dependent chromatin changes occur at OCRs unbound by Foxp3, they examined Foxp3 binding sites and chromatin accessibility among all OCRs in total Treg cells^19,21^. In contrast, we identified putative CREs and *cis* regulatory programs in single Treg cells and integrated these datasets with Treg cell-specific BATF and eTreg cell-specific Foxp3 binding profiles. This revealed that Foxp3^-^ and BATF-dependent accessibility changes occur primarily at CREs bound by both TFs in eTreg cells. Thus, our findings do not contradict previous reports, but instead reveal hidden direct and cooperative roles of Foxp3 and BATF in promoting chromatin accessibility. Notably, not all Foxp3^-^ and BATF-induced CREs were bound by both TFs. While this may indicate their indirect actions, it is also possible that they directly induce some of these CREs through transient interactions. Alternatively, their binding may be undetected due to limited sensitivity of ChIP-seq assays, particularly for CREs accessible only in subsets of eTreg cells. Mapping Foxp3 and BATF binding sites during eTreg cell differentiation and in eTreg cell subsets may reveal their direct control over such CREs.

It remains to be formally demonstrated that co-binding of BATF and Foxp3 drives increased chromatin accessibility at specific CREs operating in eTreg cells. Previous studies leveraged extensive polymorphisms in two evolutionarily divergent mouse strains to causally link TF motif disruptions to allelic imbalances in chromatin accessibility in Treg cells^19,21^. However, those authors found no significant associations between accessibility changes and disrupted BATF or FKH motifs, though AICE motifs and T*_n_*G repeats were not evaluated. This may reelect either a lack of impact by alterations in these motifs on accessibility, or a paucity of sequence variation in Foxp3 and BATF binding sites that affect accessibility, particularly among CREs. Given the likely functional importance of Foxp3^-^ and BATF-induced CREs in eTreg cells, the latter seems more plausible, as disruptions in their binding sequences would impair Treg cell functionality, reduce mouse fitness, and would therefore have been counter-selected during evolution. Establishing a causal link between Foxp3 and BATF co-binding and chromatin accessibility will require identifying their target sequences and testing whether their disruption affects accessibility—important challenges for future studies.

How do BATF and Foxp3 cooperatively promote chromatin accessibility and gene expression? While BATF is thought to act as a pioneer factor, how it induces nucleosome remodeling remains poorly understood, although recruitment of the BAF (SWI/SNF) complex has been speculated^31^. A previous study suggested that AP 1 heterodimers (FOS/JUN) promote nucleosome remodeling by recruiting the BAF complex. However, AP-1 TFs alone are insufficient. Lineage-determining TFs are also required to establish cell type-specific accessible chromatin^52^. Since Foxp3 interacts with the BAF complex^53^ and Smarcc1, one of its core subunits, is essential to promote chromatin accessibility in eTreg cells^19^, we speculate that simultaneous or sequential binding of BATF and Foxp3 facilitates nucleosome remodeling by synergistically recruiting the BAF complex. Additionally, both BATF and Foxp3 have been implicated in reorganizing chromatin architecture^31,42,48,54^. Analyses of Foxp3-deficient Treg-like cells revealed that Foxp3 is required for formation of Treg cell specific chromatin loops, particularly enhancer-promoter loops, which are closely associated with Treg cell-specific gene expression^42,54^. Interestingly, both AP-1 and FKH motifs are enriched at anchors of these loops^54^. These findings suggest that BATF and Foxp3 may mediate formation of these chromatin loops and cooperatively promote Treg cell-specific gene expression.

Our findings suggest that Foxp3 interacts with distinct TFs to direct diverse *cis* regulatory programs. We identified three categories of Foxp3-dependent programs: Foxp3-induced eTreg cell programs, Foxp3-induced cTreg/“early” eTreg cell programs, and Foxp3-repressed Tconv/cTreg cell programs. In eTreg cell programs, NR4A, NFAT, MAF, ROR, and ETS/IKZF motifs were enriched alongside AP-1 motifs, whereas NFκB and NR4A motifs were enriched in cTreg/“early” eTreg cell programs. Many members of these TF families are reported to interact with, or to be in close proximity to Foxp3 and to promote eTreg cell differentiation^32,53,55–60^. Based on these observations, we propose that Foxp3 interacts not only with BATF, but also with these TFs to cooperatively, directly, and progressively orchestrate eTreg cell programs. Consistently, our experiments revealed that Foxp3, BATF, and TCR stimulation are all necessary to promote eTreg cell differentiation, emphasizing the importance of additional TFs acting downstream of TCR signaling, such as NFκB, NR4A, and NFAT. A recent study further supports this model by suggesting that Foxp3 interacts with these TFs, including BATF, and is recruited to their target genomic sites in response to TCR signaling to regulate gene expression^60^. However, this study did not examine their roles in shaping Treg cell chromatin landscapes or their *in vivo* relevance to eTreg cell differentiation. In Foxp3-repressed Tconv/cTreg cell programs, TCF/LEF, ETS/IKZF, and RUNX motifs stood out. These TF families interact with Foxp3^53,59–62^ and participate heavily in shaping epigenetic landscapes of naïve and effector Tconv cells^63^. Thus, we suggest that Foxp3 represses these programs by directly competing with, or leveraging the repressive activity of, these TFs.

In conclusion, our study establishes Foxp3 as a master, but context-dependent regulator that interacts with other TFs, including BATF, to shape heterogeneous *cis* regulatory and transcriptional landscapes critical for functional Treg cell differentiation. Molecular dissection of these interactions will be instrumental in manipulating Treg cell function for therapeutic purposes.

## RESOURCE AVAILABILITY

### Lead contact

Requests for further information and resources should be directed to and will be fulfilled by the lead contact, Shohei Hori.

### Materials availability

The *Foxp3*^KO:hCD2^ mouse line generated in this study is available upon request and requires a Material Transfer Agreement.

## ACKNOWLEDGEMENTS

We thank Aki Minoda, Jen-Chien Chang, and Suzu Kawagoe for their invaluable advice on ATAC-seq analysis, and Kazumi Abe, Erina Ishikawa, and Yutaka Suzuki for their contributions to scMultiome analysis. We are grateful to Takako Kato, the animal and FACS facilities at RIKEN IMS and the Graduate School of Pharmaceutical Sciences, The University of Tokyo, as well as the NGS facilities at RIKEN IMS and Kazusa DNA Research Institute, for their excellent technical assistance. Supercomputing resources for bioinformatic analyses were provided by the Human Genome Center, The University of Tokyo. This work was supported by JSPS KAKENHI Grant Numbers JP18H045025, JP21H04801, JP22H05191 (to S.H.), JP17H06626, JP19K16601, JP24K10267 (to R.M.), and JP16H06279 (PAGS), by AMED under Grant Numbers JP22gm6210018, JP24bm1223012, and JP243fa627001 (to S.H.), by the Takeda Science Foundation, and by the Uehara Memorial Foundation (to SH).

## AUTHOR CONTRIBUTIONS

RM, NH, and SH designed these experiments. RM, NH, TM, ZL, SS, and YH performed all experiments. RM, TM, and SH analyzed data. RM, and HY performed bioinformatic analyses. MM, YI, WI, HK, and TK generated and provided key materials. RM and SH designed the study and wrote the manuscript.

## STAR METHODS

### EXPERIMENTAL MODEL AND STUDY PARTICIPANT DETAILS

#### Experimental animals

All mice were bred and maintained in groups of 1-5 animals per cage in specific pathogen-free facilities of the RIKEN Center for Integrative Medical Sciences and the Graduate School of Pharmaceutical Sciences, The University of Tokyo. Unless otherwise stated, males and females aged 5-20 weeks were used. All animal experiments were performed in accordance with approved protocols from the Institutional Animal Care at RIKEN and the Animal Care and Use Committee of the University of Tokyo.

B6.*Foxp3*^YFPCre^ ^35^ (JAX stock #016959), B6.*Rag1*^−/–^ ^64^ (JAX stock #002216), B6.OTII TCR transgenic ^65^ (JAX stock #004194), and B6.*Batf*^−/–^ ^66^ (JAX stock #013758) mice were purchased from The Jackson Laboratory. The following mouse strains were described previously: B6.*Foxp3*^hCD2^, B6.*Foxp3*^hCD2^ Ly5.1 ^67^, B6.*Foxp3*^R397W:hCD2^ ^26^, B6.*Foxp3*^eGFP^ ^68^, B6.*Batf*^6l^^ox^ ^69^, and B6.*Cd3e*^−/–^ ^70^ mice.

## METHOD DETAILS

### Generation of Foxp3^KO:hCD2^ allele

The *Foxp3*^KO:hCD2^ allele was generated by CRISPR-Cas9-mediated genome editing. A crRNA (Alt-R CRISPR-Cas9 crRNA; IDT) targeting exon 2 of *Foxp3* (5’-CATACCTGATGCATGAAGTG-3’) was designed, synthesized, and pre-annealed with tracrRNA (Alt-R CRISPR-Cas9 tracrRNA; IDT) at a 1:1 ratio. A mixture of crRNA/tracrRNA (1.4 μg/μl) and recombinant Cas9 protein (2.4 μg/μl; Alt-R S.p. Cas9 Nuclease V3; IDT) in Opti-MEM I (Thermo Fisher Scientieic) were used for electroporation.

Fertilized oocytes were collected following *in vitro* fertilization of superovulated oocytes and sperm, both from B6.*Foxp3*^hCD2^ mice. Superovulation was induced with equine chorionic gonadotropin (eCG) and human chorionic gonadotropin (hCG). Electroporation was performed on fertilized oocytes 6-8 hours post-insemination, followed by transfer of successfully electroporated embryos into oviducts of pseudopregnant ICR females.

To detect indel mutations in chimeric offspring, PCR amplification of genomic DNA was performed using primers flanking the edited region (left primer: 5’-GTTGCTACCGTGTGAGACTTTAGTA-3’; right primer: 5’-GATCATGGCTGGGTTGTCCAG-3’), and sequencing of the PCR product revealed the frameshift mutation described in Figure S3A, identified in one offspring, which was then mated with *Foxp3*^hCD2^ mice to establish the *Foxp3*^KO:hCD2^ line.

#### Pathological assessment

To evaluate autoimmune pathology, organs were fixed in 10% buffered formalin, and paraffin-embedded sections were stained with hematoxylin and eosin. Inflammation was graded in a blinded manner on a scale of 0 to 5, as previously described^71^.

#### Cell isolation

Single-cell suspensions of the spleen, peripheral LNs (cervical, axillar, brachial, inguinal), mesenteric LNs, and mediastinal LNs were obtained by gently forcing the tissues through a nylon mesh grid in PBS containing 2% FBS. Splenic erythrocytes were eliminated using ACK lysis buffer.

Leukocytes from the following organs were prepared as follows.

### Lung and liver

Lungs were perfused with PBS and cut into small pieces. Livers were perfused and homogenized using a steel mesh grid. Tissues were incubated in HBSS(+) containing 1% BSA, 100 U/mL collagenase type IV (Worthington), and 50 μg/mL DNase I (Worthington) for 60 min (lung) or 45 min (liver) at 37°C with occasional vortexing. Resulting suspensions were passed through a 70-μm cell strainer. Leukocytes were isolated by centrifugation at 1,000 x *g* for 20 min on a 40/70% Percoll (GE Healthcare Life Sciences) density gradient, and recovered from the interface.

### Colon

To isolate colonic lamina propria leukocytes, colons were cut longitudinally, then laterally into 1-cm pieces, thoroughly washed in PBS, and incubated twice with constant stirring in HBSS(–) containing 10 mM HEPES and 5 mM EDTA for 20 min at 37°C to remove epithelial cells. Remaining tissues were washed with PBS, cut into small pieces and digested twice with constant stirring in HBSS(+) containing 1% BSA, 200 U/mL collagenase type IV, and 100 μg/mL DNase I for 30 min at 37°C, vortexed, and passed through a 70-μm cell strainer. Leukocytes were isolated by centrifugation at 1,000 x *g* for 20 min on a 40/80% Percoll density gradient and recovered from the interface.

### Skin

Ears were excised, separated into ventral and dorsal sheets, and incubated in RPMI-1640 medium containing 10% FBS, 320 U/mL collagenase type IV, and 100 μg/mL DNase I for 90 min at 37°C in 5% CO_2_. Digested tissues were then homogenized using a nylon mesh grid and passed through a 70-μm cell strainer to obtain single-cell suspensions.

#### Flow cytometry

Cells were pre-incubated with anti-FcγR mAb (clone 2.4G2, prepared in-house) to block Fc receptor binding before staining with appropriate mAbs at 4°C for 30 min. For detecting CCR7 expression, cells were incubated with biotinylated anti CCR7 mAb (diluted 1:10) at 4°C for 60 min, followed by staining with PE^-^ or PE CF594-conjugated streptavidin. All staining was done in PBS containing 2% calf serum and 0.02% sodium azide. To exclude dead cells, cells were suspended in staining buffer containing 0.5 μg/mL propidium iodide (Sigma-Aldrich) or 1 μM SYTOX Blue (Thermo Fisher Scientieic) before FCM analysis.

For intracellular staining of transcription factors and CTLA-4, cells were first stained for surface antigens, followed by staining with Fixable Viability Dye eFluor 780 (Thermo Fisher Scientieic). Cells were then fixed, permeabilized, and stained for intracellular antigens using the Foxp3/Transcription Factor Staining Buffer Set (Thermo Fisher Scientieic) according to the manufacturer’s instructions. Unconjugated anti-BATF rabbit mAb (D7C5, Cell Signaling Technology) was revealed using PE F(ab’)_2_ donkey anti-rabbit IgG (Thermo Fisher Scientieic) in the presence of mouse IgG, rat IgG, Syrian hamster IgG (Jackson ImmunoResearch or Sigma-Aldrich) to block non-specific binding.

For intracellular cytokine staining, cells were first stained for surface antigen and with Fixable Viability Dye eFluor 780, fixed with PBS containing 4% paraformaldehyde for 10 min at room temperature, permeabilized, and stained for cytokines using Perm/Wash Buffer (BD Biosciences).

Flow cytometric analysis was performed on FACSCanto II, FACSAria III (BD Biosciences), or CytoFLEX S (Beckman Coulter), and data were analyzed using FlowJo software (Tree Star, Inc.).

#### Cell sorting

Cell sorting was performed using FACSAria II or III cell sorters (BD Bioscience). Spleen and LN cells were prepared in PBS containing 2% FBS. After depletion of erythrocytes with ACK lysis buffer, spleen cells were pooled with LN cells and depleted of B220^+^, CD8β^+^, CD11b^+^, Gr1^+^, and adherent cells by panning. To sort CD4^+^ T cell subsets from mice with a *Foxp3*^hCD2^ reporter allele, cells were stained with PE or APC-conjugated anti-human CD2, along with other fluorescent mAbs. After labeling with anti-PE or APC Microbeads (Miltenyi Biotec), cells were separated into hCD2^+^ and hCD2^-^ cells on MS or LS columns before being subjected to FACS sorting. To sort OTII Tn cells, pooled spleen and LN cells were stained with fluorescent mAbs and sorted into CD44^low^CD62L^high^hCD2^-^CD4^+^ cells without pre-enrichment by panning and MACS. Purity of sorted cells was consistently >98 %.

#### RV vectors and production of RV supernatants

Mouse Foxp3 cDNA cloned in MSCV-IRES-GFP (MIGR1) ^5^ and mouse BATF cDNA cloned in MSCV-IRES-Thy1.1 retroviral vectors^26^ were previously described. Plat-E packaging cells^72^ were cultured in DMEM (high glucose) (Sigma-Aldrich), supplemented with 10% FBS and 10 mM HEPES, and transfected with retroviral vectors using FuGENE 6 Transfection Reagent (Promega). Viral supernatants were collected 48 hr after transfection.

#### In vitro T cell activation and retroviral transduction

T cells were cultured in RPMI-1640 medium supplemented with 10% FBS (Hyclone or Gibco), GlutaMAX^TM^ (Thermo Fisher Scientieic) or L-alanyl-L-glutamine (Nacalai), 10 mM HEPES (Sigma-Aldrich), 1 mM sodium pyruvate (Sigma-Aldrich), 100 U/mL penicillin, 100 μg/mL streptomycin (Sigma-Aldrich), 50 μM 2-ME. For retroviral transduction, T cells (1.0 x 10^6^ cells/mL) were stimulated with Dynabeads Mouse T-Activator CD3/CD28 (Thermo Fisher Scientieic) (1.0 x 10^6^/mL) in the presence of 10 ng/mL recombinant mouse (rm) IL-2 (R&D Systems) in 48-well plates. Twenty-four hours later, cells were transduced using an equal volume of retroviral supernatant and 6 μg/mL polybrene (Sigma-Aldrich) and centrifuged at 1,580 x *g* for 60 min at 32°C. The culture medium was replaced 12-16 hr after transduction, and cells were maintained and expanded in medium containing 10 ng/mL rmIL-2. On day 4 after the initial stimulation, cells were harvested for adoptive transfer experiments.

#### Adoptive transfer and intranasal OVA administration

Donor cells were labeled with 10 μM CPD eFluor 450 (Thermo Fisher Scientieic), suspended in sterile PBS, and injected intravenously into sex-matched recipient mice. Sublethally irradiated (6 Gy) Ly5.1 or Ly5.1/5.2 *Foxp3*^hCD2^ mice were used as recipients of retrovirally transduced hCD2^+^CD4^+^ T cells from Ly5.2 *Foxp3*^R397W:hCD2^*Batf*^−/–^ mice (5.0 x 10^5^ cells/recipient). After irradiation, recipient mice were provided with drinking water containing 0.5 mg/mL neomycin sulfate (Sigma-Aldrich) and 100 U/mL polymyxin B sulfate. For retrovirally transduced Ly5.2 OTII CD4^+^ T cell transfers, Ly5.1 *Foxp3*^hCD2^ mice were used as recipients (1.0 x 10^5^ cells/recipient). These mice received 4 ∝g OVA protein (Sigma-Aldrich) or sterile PBS via intranasal administration on day 1 post-transfer.

### Cytokine assay and in vitro suppression assay

#### Cytokine assay

Cells were stimulated for 4 hr at 37℃ with 50 ng/mL PMA and 500 ng/mL ionomycin in the presence of GolgiStop (BD Biosciences) before intracellular staining.

#### *In vitro* suppression assay

hCD2^+^CD4^+^ T cells from Ly5.2 *Batf*^−/–^ *Foxp3*^R397W:hCD2^ or *Batf*^−/–^ *Foxp3*^hCD2^ mice were retrovirally transduced with GFP RV and Thy1.1 RV. On day 3 after transduction, GFP^+^Thy1.1^+^hCD2^+^CD4^+^ cells were sorted and used as Treg cells. MACS-sorted hCD2-CD4^+^ cells from Ly5.1 *Foxp3*^hCD2^ mice were labeled with 10 μM CPD eFluor 450 and used as responder T cells. Responder T cells (5.0 x 10^4^ cells/well) were cultured alone or together with a titrated number of Treg cells in the presence of 0.5 μg/mL anti-CD3ε mAb (145-2C11) and X-irradiated (20 Gy) *Cd3e*^−/–^ splenocytes as antigen-presenting cells (2.0 x 10^5^ cells/well) in 96 well U bottom plates. On day 3, cells were stained for Ly5.1, Ly5.2, and CD4 and analyzed by FCM.

#### Bulk RNA-seq

The following cell populations were sorted from pooled spleen and LN cells and subjected to RNA-seq analysis (*n*=2 per group): YFP^+^hCD2-CD4^+^ cells from *Batf*^6l^^ox/6l^^ox^ *Foxp3*^YFPCre/hCD2^ mice (BATF cKO Treg) and *Batf*^+/+^ *Foxp3*^YFPCre/hCD2^ mice; hCD2^+^CD4^+^ (total Treg) and hCD2-CD4^+^ (total Tconv) cells from *Foxp3*^hCD2^ mice; hCD2^+^CD4^+^ cells from *Batf*^−/–^ *Foxp3*^hCD2^ mice (BATF KO Treg); hCD2^+^CD4^+^ cells from *Foxp3*^R397W:hCD2/GFP^ (R397W Treg) and *Foxp3*^KO:hCD2/GFP^ (Foxp3 KO Treg) mice, and their respective *Foxp3*^WT:hCD2/GFP^ control mice; CD44^low^CD62L^high^CCR7^high^ hCD2^+^CD4^+^ (cTreg), CD44^high^CD62L^low^CCR7^low^ hCD2^+^CD4^+^ (eTreg), CD44^low^CD62L^high^CCR7^high^ hCD2-CD4^+^ (Tn), and CD44^high^CD62L^low^CCR7^low^ hCD2^-^ CD4^+^ (Teff) cells from *Foxp3*^hCD2^ mice.

Total RNA was isolated from sorted cells using Isogen (Nippon Gene) and quantified using the Quant-iT^TM^ RiboGreen RNA Assay Kit (Thermo Fisher Scientieic) or the QuantiFluor RNA system (Promega). RNA quality was assessed on a 2100 Agilent Bioanalyzer. RNA-seq libraries were prepared from 50-300 ng total RNA using either the NEBNext Ultra Directional RNA Library Prep Kit for Illumina or the NEBNext Ultra II RNA Library Prep Kit for Illumina (New England Biolabs) with 14 or 15 cycles of PCR amplification. Library concentration was quantified using a KAPA Library Quantification Kit (Kapa Biosystems). Barcoded libraries were sequenced as single-end 50-bp reads on a Hiseq 1500 (Illumina) or paired-end 100-bp reads on a DNBSEQ-G400 (MGI).

#### Bulk ATAC-seq

ATAC-seq libraries of the following populations sorted from pooled spleen and LN cells were prepared according to the standard ATAC-seq protocol^73^ (*n* = 2 per group): CD44^low^CD62L^high^CCR7^high^ hCD2^+^CD4^+^ (cTreg), CD44^high^CD62L^low^CCR7^low^ hCD2^+^CD4^+^ (eTreg), CD44^low^CD62L^high^CCR7^high^ hCD2-CD4^+^ (Tn), CD44^high^CD62L^low^CCR7^low^ hCD2-CD4^+^ (Teff), and hCD2^+^CD4^+^ (WT Treg) cells from *Foxp3*^hCD2^ mice, and hCD2^+^CD4^+^ (BATF KO Treg) cells from *Batf*^−/–^ *Foxp3*^hCD2^ mice. Sorted 5 x 10^4^ cells were washed with PBS, treated with cold lysis buffer (10 mM Tris-HCl, pH7.4; 10 mM NaCl; 3 mM MgCl_2_; 0.1 % IGEPAL CA-630), reacted with Tagment DNA TDE1 Enzyme (Illumina), and amplified by PCR for 9-11 cycles.

ATAC-seq libraries of hCD2^+^CD4^+^ T cells sorted from pooled spleen and LN cells of *Foxp3*^R397W:hCD2/GFP^, *Foxp3*^KO:hCD2/GFP^, and their respective *Foxp3*^WT:hCD2/GFP^ control mice were prepared according to the omni-ATAC-seq protocol^74^ (*n* = 2 per group). Sorted 5 x 10^4^ cells were washed with PBS, treated with lysis buffer (10 mM Tris HCl, pH7.4; 10 mM NaCl; 3 mM MgCl_2_; 0.1 % NP-40; 0.1% Tween-20; 0.01% Digitonin), reacted with Tn5 transposase (Diagenode), and amplified by PCR for 12 cycles. Library concentration was quantified using the KAPA Library Quantification Kit (Kapa Biosystems). Barcoded libraries were sequenced as single-end 50-bp reads on a Hiseq 2500 (Illumina) or paired-end 100-bp reads on a DNBSEQ-G400 (MGI).

#### scATAC+RNA-seq (scMultiome)

WT Tconv (hCD2-CD4^+^) and WT Treg (hCD2^+^CD4^+^) cells from *Foxp3*^hCD2^ mice, BATF KO Treg cells from *Batf*^−/–^ *Foxp3*^hCD2^ mice, and R397W Treg cells from *Foxp3*^R397W:hCD2/GFP^ mice, all sorted from pooled spleens and LNs, were subjected to scMultiome analysis (*n* = 2 per group). Sorted cells were washed with PBS containing 0.04% BSA, treated with lysis buffer (10 mM Tris-HCl, pH7.4; 10 mM NaCl; 3 mM MgCl_2_; 0.1% Tween-20; 0.1% NP-40; 0.01% digitonin; 1% BSA; 1 mM DTT; 1 U/μL RNase inhibitor) for 3 min on ice, and washed three times with wash buffer (10 mM Tris-HCl, pH7.4; 10 mM NaCl; 3 mM MgCl_2_; 1% BSA; 0.1% Tween-20; 1 mM DTT; 1 U/μL RNase inhibitor) to collect nuclei. Isolated nuclei were passed through 40-μm cell strainers (Falcon) and counted using automated cell counters (Thermo Fisher Scientieic).

A total of 5-10 x 10^3^ isolated nuclei were subjected to scMultiome library preparation using the Chromium Next GEM Single Cell Multiome ATAC + Gene Expression Reagent (10x Genomics) according to the manufacturer’s instructions. Libraries were sequenced on a NovaSeq 6000 (Illumina).

#### ChIP-seq

BATF ChIP-seq was performed on hCD2^+^CD4^+^ and hCD2-CD4^+^ cells from *Foxp3*^hCD2^ mice (*n* = 2 per group). Foxp3 ChIP-seq was conducted on hCD2^+^CD4^+^ (WT Treg), CD44^low^CD62L^high^CCR7^high^ hCD2^+^CD4^+^ (cTreg), CD44^high^CD62L^low^CCR7^low^ hCD2^+^CD4^+^ (eTreg) cells from *Foxp3*^hCD2^ mice, and hCD2^+^CD4^+^ cells from *Batf*^−/–^

*Foxp3*^hCD2^ mice (BATF KO Treg) (*n* = 2 per group). A total of 5-12 x10^6^ sorted cells were fixed with 1% formaldehyde for 30 min at room temperature, lysed sequentially with lysis buffer 1 (50 mM HEPES, pH7.5; 140 mM NaCl; 1 mM EDTA; 10% glycerol; 0.5% NP-40; 0.25% Triton X-100; 5 mM DTT; protease inhibitors (Roche)), lysis buffer 2 (10 mM Tris-HCl, pH8.0; 200 mM NaCl; 1 mM EDTA; 0.5 mM EGTA; protease inhibitors), and lysis buffer 3 (10 mM Tris-HCl, pH8.0; 300 mM NaCl; 1 mM EDTA; 0.5 mM EGTA; 0.1% sodium deoxycholate; 0.5% N-lauroylsacrosine; protease inhibitors), followed by sonication.

Cell extracts were incubated overnight at 4℃ with 6.5 μg/mL of anti-BATF mAb (clone: D7C5, Cell Signaling Technology) or 14 μg/mL affinity-purified polyclonal anti-Foxp3 antibodies^26^. Dynabeads Protein G (Dynal) were added for 4 hr at 4℃. Dynabeads were washed successively with low salt buffer (20 mM Tris-HCl, pH8.0; 150 mM NaCl; 0.1% SDS; 1% Triton-X100; 2 mM EDTA), high salt buffer (20 mM Tris-HCl, pH8.0; 500 mM NaCl; 0.1% SDS; 1% Triton-X100; 2 mM EDTA), RIPA buffer (50 mM HEPES, pH7.6; 500 mM LiCl; 1% NP-40; 0.7% sodium deoxycholate; 1 mM EDTA), and TE buffer containing 50 mM NaCl. Elution was performed by heating the Dynabeads in elution buffer (50 mM Tris-HCl, pH8.0; 1% SDS; 10 mM EDTA) for 10 min at 65℃ with vortexing, and crosslinking was reversed by overnight incubation at 65℃.

Immunoprecipitated DNA and whole-cell extract input DNA were treated with RNase A and proteinase K, purified using a MinElute PCR Purification Kit (QIAGEN), and processed for ChIP-seq library preparation. Libraries were prepared using the NEBNext ChIP-Seq Library Prep Reagent Set for Illumina or the NEBNext Ultra II DNA Library Prep Kit for Illumina (New England Biolabs) according to manufacturer instructions and amplified by PCR for 15 cycles. A KAPA Library Quantification Kit (Kapa Biosystems) was used to quantify library concentration. Barcoded libraries were sequenced as single-end 50-bp reads on a Hiseq 1500 (Illumina).

#### Co-immunoprecipitation and western blotting

hCD2^+^ and hCD2^-^ CD4^+^ T cells from *Foxp3*^hCD2^ mice (1.0 x 10^6^ cells/well) were stimulated with or without an equal number of Dynabeads Mouse T-Activator CD3/CD28 (Thermo Fisher Scientieic) in the presence of 10 ng/mL rmIL-2 in 48-well plates (1 mL/well). On day 3, Dynabeads were removed, and nuclear and cytosolic fractions were isolated from 5.0-10 x 10^6^ cells using the Nuclear Complex Co-IP Kit (Active Motif) according to manufacturer instructions. A portion of the nuclear fraction was reserved as input. The remaining nuclear fraction was diluted with IP low-incubation buffer (Active Motif), incubated with anti-BATF mAb (D7C5, Cell Signaling Technology) at 1:100 dilution overnight at 4℃, and further incubated with 0.8 mg/mL of FG beads Protein G (TAMAGAWA SEIKI) for 3 hr at 4℃ with constant rotation. Following washing with IP low wash buffer, bound proteins were eluted in Laemmli sample buffer by heating at 98℃ for 10 min, separated by SDS-PAGE alongside input and cytosolic samples, and transferred to Immobilon-P PVDF membranes (Millipore).

Membranes were blocked with Tris-Buffered Saline containing 5% skim milk and 0.2% Tween-20 for 30 min, then incubated with anti-Foxp3 (eBio7979, Thermo Fisher Scientieic), anti-BATF (D7C5, Cell Signaling Technology), anti-Lamin B (M-20, Santa Cruz), or anti-β actin (AC-74, Sigma) diluted in Can Get Signal Solution 1 (Toyobo) overnight. After washing, membranes were incubated with HRP conjugated TrueBlot ULTRA anti-mouse IgG (Thermo Fisher Scientieic), TrueBlot anti-rabbit IgG (Rockland), or TrueBlot anti-Goat IgG (Rockland) secondary antibodies diluted in Can Get Signal Solution 2 for 3 hr. Signals were detected using Chemi-Lumi One Ultra (Nacalai).

## QUANTIFICATION AND STATISTICAL ANALYSIS

### Data analysis for bulk RNA-seq

Sequence reads were aligned to the mouse reference genome (UCSC mm10) and raw read counts were calculated for each gene using *STAR*^75^ (version 2.6.1c). For gene expression comparison among multiple groups, the likelihood ratio test implemented in *edgeR*^76^ (version 3.38.4) was applied, defining differentially expressed genes (DEGs) as those with an FDR ≤0.05. For pairwise comparisons, Fisher’s exact test in *edgeR* was used, with DEGs defined as genes with an FDR ≤0.05 and an absolute log2 fold-change (∣log2FC∣) ≥0.5. Non-differential expression was denied by FDR >0.05 or ∣log2FC∣ <0.5.

Hierarchical clustering and heatmap generation were performed using the *seaborn clustermap* function in *Python* (version 3.6.4), employing the Euclidean metric for distance calculations. Over-representation analysis^77^ of BATF-induced genes and genes with TSSs nearest to BATF-induced OCRs in the Molecular Signatures Database (MSigDB)^78,79^ (msigdb7.5.1.entrez.gmt) was conducted using *clusterProMiler*^80^ (version 4.4.4).

### Data analysis for bulk ATAC-seq

Sequence reads were aligned to the mouse reference genome (UCSC mm10) using *Bowtie2*^81^ (version 2.4.1). Peak calling was performed using *MACS2*^82,83^ (version 2.2.7.1) with a *q*-value threshold of 0.01. Identification of consensus OCRs and differentially accessible OCRs was performed using *DIFFBIND*^84,85^ (version 3.0.15). Blacklist regions were excluded, and consensus OCRs were defined as regions detected in both biological replicates of at least one cellular group. Differentially accessible OCRs were identified as regions with *q*-values ≤ 0.05. Gene annotation for ATAC-seq peaks was conducted using the *annotatePeaks.pl* function in *HOMER*^86^ (version 4.9.1). Motif analysis of OCRs was performed using *MEME ChIP*^87^ (version 5.5.5) in differential enrichment mode.

### Data analysis for scMultiome

#### Preprocessing and UMAP visualization

Alignment, filtering, and barcode counting were performed using *Cell Ranger ARC* (version 2.0.0, 10x Genomics) with the mm10 reference genome provided, generating cell vs. gene matrices of UMI counts for scRNA-seq data and cell vs. fragment matrices for scATAC-seq data. Downstream analysis was performed using *Seurat*^88^ (version 4.3.0, 5.0.3 or 5.1.0) and *Signac*^40^ (version 1.9.0, 1.12.0 or 1.13.0). Low-quality cells were excluded based on the following criteria: percent mitochondria ≥10, TSS.enrichment ≤3, nCount_RNA ≤2,000 or ≥20,000, nCount_ATAC ≤5,000 or ≥50,000, blacklist ratio ≥0.05, and fraction of reads in peaks ≤0.4.

For scATAC-seq data, peak calling was performed using *MACS2* (v2.2.7.1). Peaks in non-standard chromosomes and genomic blacklist regions were filtered out using *keepStandardChromosomes* and *subsetByOverlaps* functions, respectively. Fragment counts for resulting peaks were quantified using the *FeatureMatrix* function in *Signac*, and assigned to a new assay, which was used for downstream analysis. Fragment counts were normalized using the *RunTFIDF* function (method = 1). After feature selection using *FindTopFeatures* (min.cutoff = 5), dimension reduction was carried out using *RunSVD*. For scRNA-seq data, normalization and scaling of UMI counts were performed using *NormalizeData* and *ScaleData*, respectively. Highly variable genes were identified using the *FindVariableFeatures* function (selection.method = “vst” and nfeatures = 2000). PCA of count data of resulting variable genes was conducted using the *RunPCA* function (npcs = 30). Datasets from two independent experiments for both scATAC-seq and scRNA-seq were integrated using the *RunHarmony* function of *harmony*^89^ (version 1.2.0) based on normalized counts.

For scATAC-seq data, dimensionality reduction and cluster identieication were performed using *RunUMAP* (dims = 2:50), *FindNeighbors* (dim = 2:30), and *FindClusters* (algorithm = 3, resolution = 0.5), resulting in the UMAP plot shown in Figure S5B. Clusters 0-4 and 6-8 were extracted for re-clustering. Dimensionality reduction and cluster identification were repeated using *FindTopFeatures* (min.cutoff = 5), *RunSVD*, *RunUMAP* (dims = 2:50), *FindNeighbors* (dim = 2:30), and *FindClusters* (algorithm = 3, resolution = 0.5), resulting in the UMAP plot shown in Figure 4A. Clusters 3 and 5-7 were further re-clustered, with dimensionality reduction and cluster identification performed using *FindTopFeatures* (min.cutoff = 5), *RunSVD*, *RunUMAP* (dims = 2:50), *FindNeighbors* (dim =2:30), and *FindClusters* (algorithm =3, resolution = 0.3), yielding the UMAP plot shown in Figure 4F.

#### Calculation and visualization of gene expression scores and accessibility scores

The *AddModuleScore* function in *Seurat* was used to calculate gene expression scores for gene signatures of pTreg cells induced by commensal bacteria (kindly provided by Dr. Alexander Rudensky^37^) and colonic RORγt^+^ pTreg cells^36^ in single cells. For the latter, the public microarray dataset GSE68009, available on the Gene Expression Omnibus, was analyzed as follows: probe counts were obtained from CEL files and normalized using *oligo*^90^ (version 1.60.0), with gene annotation added for each probe using *affycoretools* (version 1.68.1) and *mogene10sttranscriptcluster.db* (version 8.8.0). Gene expression comparisons were performed using *lmFit* and *eBayes* functions in *limma*^91^ (version 3.52.4). Genes upregulated in colonic RORγt^+^ compared to RORγt^-^ Treg cells (*p* ≤ 0.05 & log2 fold change ≥ 0.5) were defined as the colonic RORγt^+^ pTreg cell gene signature. Accessibility scores for specific sets of OCRs in single cells were calculated using the *AddChromatinModule* function in *Signac*. These scores were visualized on UMAPs using the *FeaturePlot* function (max.cutoff = q95) and as violin plots using the *VlnPlot* function in Figures 4D, 4G, 5B, 5C, 6C, and S6E. For Figures 6D and 6E, accessibility scores for CREs associated with each topic were averaged for each cluster or subcluster. OCRs that differed in accessibility between WT Treg cells vs. BATF KO or R397W Treg cells were identified using the *FindMarkers* function (min.pct = 0.01, test.use = LR and logfc.threshold = 0.1) and defined as those with FDR ≤ 0.05.

#### Identification of variable TF motifs and calculation of TF motif activity scores

The *computeVariability* function in *chromVAR*^38^ (version 1.16.0, 1.24.0) was used to calculate variability scores for TF motifs from the JASPAR 2022 database (version 0.99.7) in single cells. The 100 TF motifs with the highest variability scores were selected for analysis in Figure 5D, 5E, and 6H.

*ChromVAR* was also used to measure changes in chromatin accessibility at ATAC seq peaks containing specific TF binding motifs across single cells, enabling calculation of per-cell TF motif activity scores. Motif annotations were assigned to each peak using the *AddMotifs* function in *Signac*, incorporating position frequency matrices from JASPAR 2022 (version 0.99.7) or position weight matrices of *homer_pwms* in *chromVARmotifs* (version 0.2.0). The *RunChromVAR* function in *Signac* was subsequently used to calculate TF motif activity scores in single cells. In Figure 5D and 5E, these scores were averaged per cluster or subcluster and represented as heatmaps with hierarchical clustering. In Figure 5F and 5G, motif scores were visualized on UMAPs using the *FeaturePlot* function (max.cutoff = q95) and as violin plots using the *VlnPlot* function. BATF (MA1634.1) and Foxp3 (MA0850.1) motifs were sourced from JASPAR 2022, while the AICE motif (bZIP:IRF(bZIP,IRF)/Th17-BatF-ChIP-Seq(GSE39756)/Homer) was sourced from *homer_pwms*.

#### Identification and characterization of OCR-gene links

To identify OCR-gene links, we considered all ATAC-seq peaks within ±500 kb of the TSS for each gene. The *LinkPeaks* function in *Signac* was used to compute Pearson correlation coefficient and *p*-value between gene expression levels and ATAC peak counts for each OCR-gene pair. OCR-gene pairs with *p*-values of ≤0.05 and absolute Pearson correlation coefficient ≥0.05 were retained and considered OCR-gene links. Visualization of these links and aggregated (pseudo-bulk) genome coverage tracks was performed using the *Coverageplot* function.

In Figure 6A and S6C, Treg and Tconv cell SE genes, as defined by H3K27ac ChIP seq^41^, were used. Additionally, enhancer-promoter loops detected in Treg and/or Tonv cells, as defined by H3K27ac HiChIP^42^, were utilized in Figure S6D.

#### Topic modeling on scATAC-seq data of CREs

To model *cis*-regulatory programs, we performed topic modeling using *cisTopic*^43^ (version 0.3.0), focusing on linked OCRs (CREs) rather than all OCRs. Input data consisted of a cell-by-count matrix of CREs extracted from a *Seurat* object, in which each count represented the accessibility of a CRE in individual cells. The *runWarpLDAModels* function in *cisTopic* was used with 500 iterations for each model. Several candidate topic numbers (2, 6, 8, 10, 12, 14, 16, 20, 30, and 40) were tested to determine the optimal topic number. Model selection was performed using the default ‘derivative’ method, which evaluates changes in log-likelihood across different topic numbers. A model with 16 topics was selected, as it provided a sufficient fit to both chromatin accessibility patterns and single-cell data, while increasing the topic number, i.e., to 20, did not significantly improve the log likelihood. To identify representative OCRs for each topic, the *binarizecisTopics* function in *cisTopic* was applied with a probability threshold of 0.9 (thrP = 0.9).

#### Motif identieication in OCRs

*FIMO*^92^ (version 5.5.5) was used to identify motifs in CREs or all OCRs. The RYMAAYA motif was identified using a default *p*-value cutoff of 1 x 10-^4^, while (T*_n_*G)_10_ (*n*=2–5) repeats were identified with a *q*-value cutoff of 0.01 for multiple testing correction in long sequences. The *matchMotifs* function in *Signac* was applied with a default *p*-value cutoff of 5 x 10^-5^ to identify the Foxp3 motif (MA0850.1), BATF motif (MA1634.1), and the top 100 most variable TF motifs identified earlier, all derived from JASPAR 2022. Additionally, the AICE motif (bZIP:IRF(bZIP,IRF)/Th17-BatF-ChIP-Seq(GSE39756)/Homer) was identified using *homer_pwm* from *HOMER*^86^ (version 4.9.1).

### Data analysis for ChIP-seq

Sequence reads were aligned to the mouse reference genome (UCSC mm10) using *Bowtie2* (version 2.4.1). Peak calling was performed with *MACS2* (version 2.2.7.1), using input chromatin as control and a *q*-value threshold of 1 x 10-^4^. Identification of consensus Foxp3 or BATF binding sites and differential binding analysis were conducted using *DIFFBIND* (version 3.0.15, 3.12.0). Blacklist regions and greylist regions (*p*-value < 0.999) were excluded. Consensus Foxp3 or BATF binding sites were defined as regions detected in both biological replicates of at least one cellular group, while differential binding sites were defined as regions with *q* values ≤ 0.05.

ChIP-seq peaks were scaled with *genomeCoverageBed* function in *bedtools*^93^ (version 2.27.1, 2.31.1), converted to bigwig format with *ucsc-bedgraphtobigwig*^94^ (version 377), and visualized with *IGV*^95^ (version 2.16.2). Overlaps between ChIP seq and scATAC-seq peaks were analyzed using the *intersect* function of *bedtools* (version 2.27.1, 2.31.1).

Motif analysis of Foxp3 and BATF binding sites was performed with *MEME ChIP*^96^ (version 5.5.5). For Figure 7D, differential enrichment mode was employed to identify motifs that are enriched in binding sites of interest relative to the remaining binding sites, using sequences of binding sites of interest as primary sequences and sequences of the remaining binding sites as control sequences. For Figure S7A and S7B, the classic mode was used to identify motifs enriched in the provided binding sites relative to a random model, based on frequencies of letters in these sequences.

### Other statistical analysis

Statistical analyses were performed using *Prism* (versions 6 and 9, GraphPad Software), *R* (version 4.0.2, 4.1.0, 4.2.1 or 4.3.2), or *Python* (version 3.6.4). Specific statistical tests used to evaluate significance are detailed in figure legends. All error bars denote mean ± SD. Differences with *p*-values ≤0.05 were considered statistically significant. Symbols used are as follows: n.s., not significant; * *p* ≤0.05; ** *p* ≤0.01; *** *p* ≤0.001; **** *p* ≤0.0001. *n* represents the number of biological replicates.

## SUPPLEMENTAL FIGURE LEGENDS

**Figure S1:**
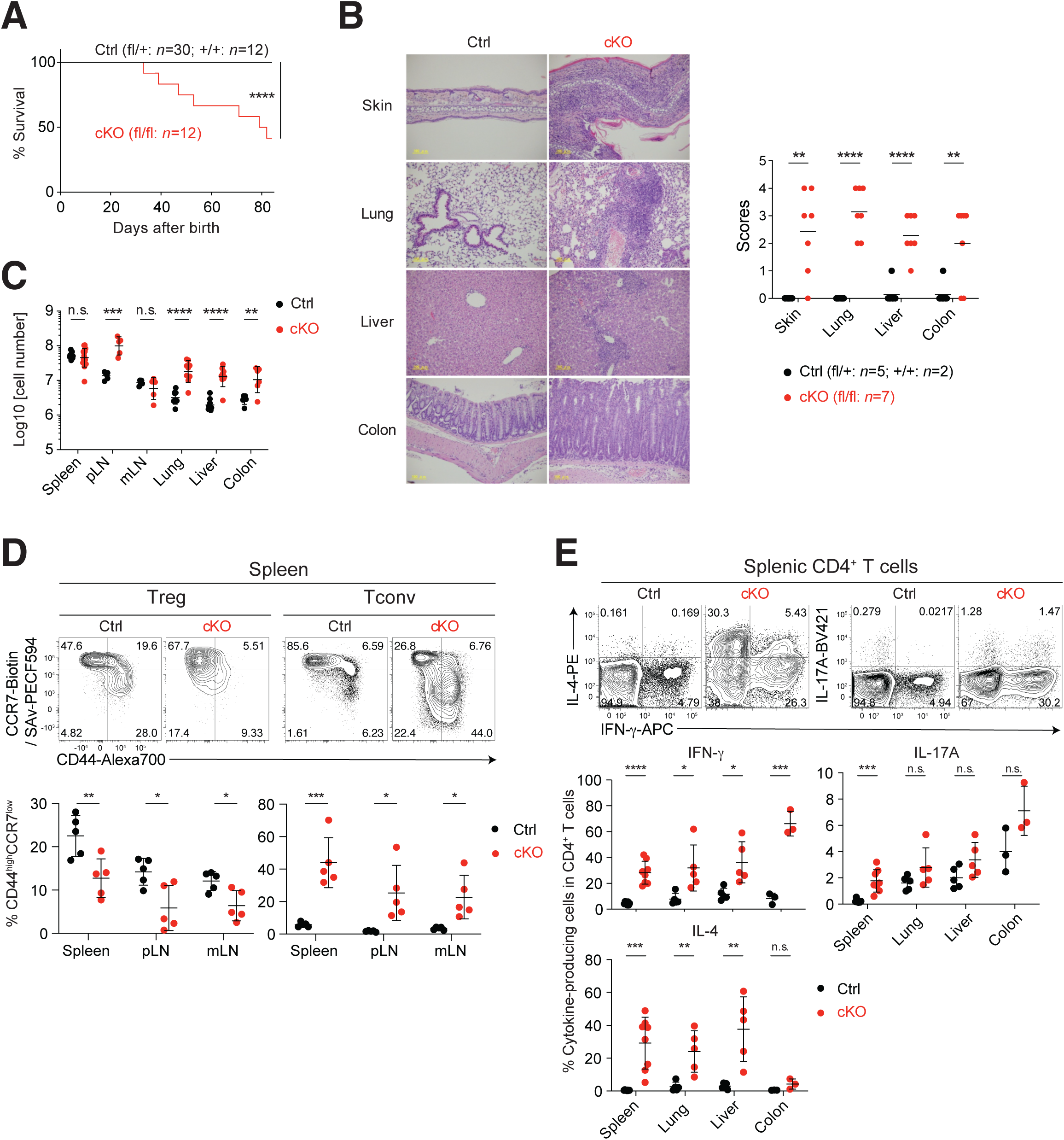
Fatal multi-organ inflammation in Treg cell-specific BATF-deficient mice, related to Figure 1. (A) Survival of cKO (*Batf*^6l/6l^ *Foxp3*^YFPCre/Y^ or *Foxp3*^YFPCre/YFPCre^) and control (Ctrl; *Batf*^6l/+^ or *Batf*^+/+^ *Foxp3*^YFPCre/Y^ or *Foxp3*^YFPCre/YFPCre^) mice. (B) Histological analysis of indicated organs by hematoxylin and eosin staining. Inflammation scores are summarized. (C) Numbers of leukocytes isolated from indicated organs (*n*=6–10). (D) FCM of Foxp3^+^ (Treg) and Foxp3^-^ (Tconv) CD4^+^TCRβ^+^ cells. Percentages of CD44^high^CCR7^low^ cells are summarized (*n*=5). (E) FCM of CD4^+^TCRβ^+^ cells. Percentages of cytokine-producing cells among CD4^+^TCRβ^+^ cells are summarized (*n*=3–8). Mice aged 5–6 (D) or 5–8 (B, C, E) weeks were analyzed. Each dot represents an individual mouse. Data analyzed by the log-rank test (A) or two-way ANOVA with Sidak’s multiple comparisons test (B–E).

**Figure S2:**
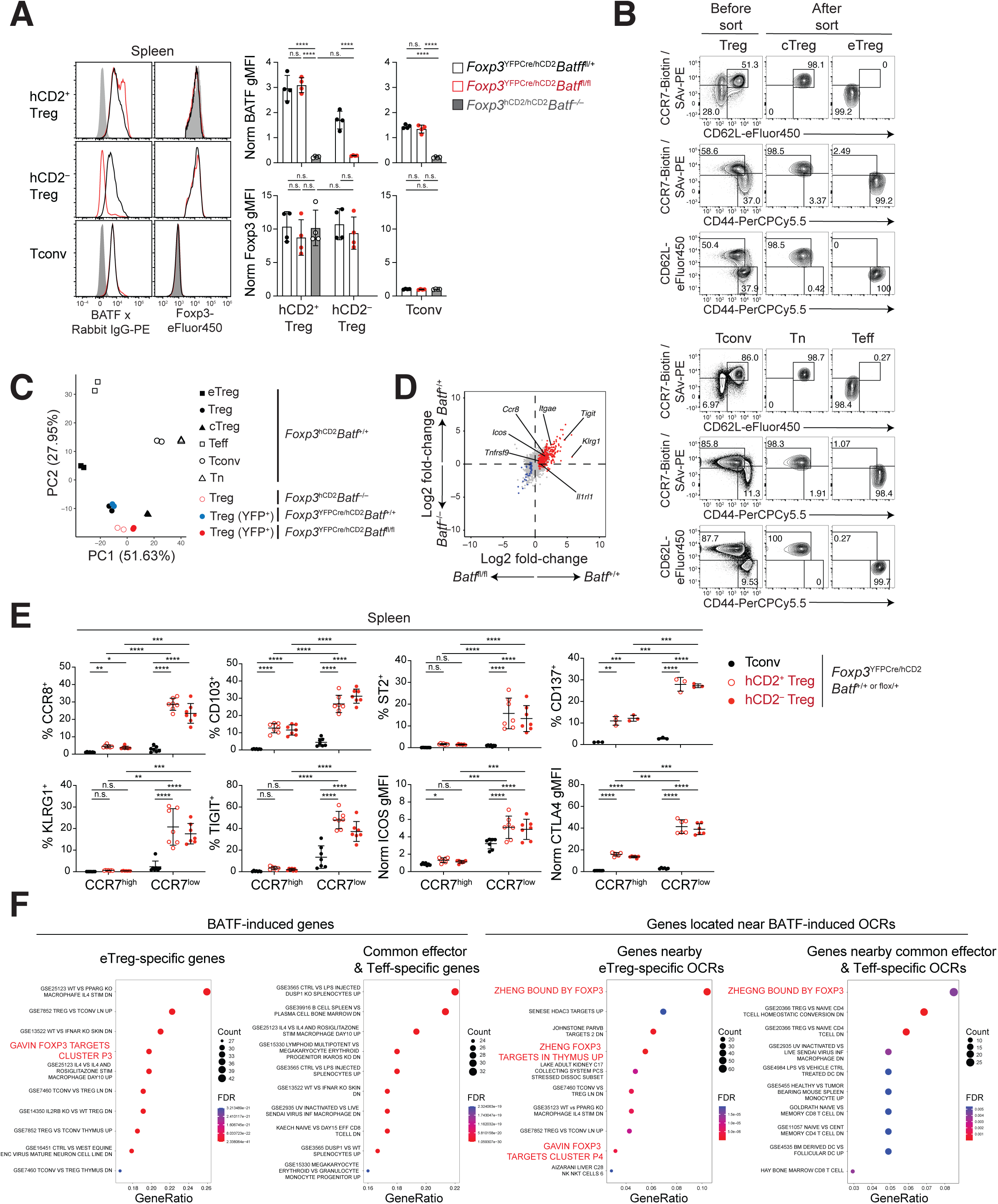
Characterization of BATF-deficient Treg cells, related to Figure 1. (A) FCM of CD4^+^ T cell subsets from indicated mice. gMFI values of BATF and Foxp3 are normalized to mean values of Tconv cells in each experiment and summarized in graphs (*n*=4). (B) Sorting of cTreg, eTreg, Tn, and Teff cells from pooled spleens and LNs of *Foxp3*^hCD2^ mice. FCM of Treg and Tconv cell subsets before and after sorting. Sorted cells were subjected to RNA-seq or ATAC-seq analysis (*n*=2). (C) PCA of RNA-seq data using the 1,000 most variable genes with the lowest FDR by the likelihood ratio test. (D) Comparison of gene expression between BATF cKO and BATF KO Treg cells. hCD2^+^CD4^+^ Treg cells from pooled spleens and LNs of *Batf*^−/–^ or *Batf*^+/+^ *Foxp3*^hCD2^ mice were subjected to RNA-seq analysis (*n*=2). FC vs. FC plot comparing the effects of Treg cell-specific BATF deficiency vs. germline BATF deficiency on gene expression. BATF-induced (red) and BATF-repressed (blue) (defined in Figure 1B) genes are highlighted. (E) Percentages of cells expressing indicated molecules or gMFI of ICOS or CTLA-4 in indicated subsets (*n*=3–7). gMFI values are normalized to mean values in Tconv cells in each experiment. (F) Over-representation analysis of BATF-induced genes and genes near BATF induced OCRs within the Molecular Signatures Database (MSigDB) gene sets. ‘Count’ refers to the number of genes in the MSigDB gene set, and ‘GeneRatio’ represents the ratio of genes in the MSigDB gene set to the analyzed gene set. Data analyzed by one-way ANOVA with Tukey’s multiple comparisons test (A) or two-way ANOVA with Sidak’s multiple comparisons test (E).

**Figure S3:**
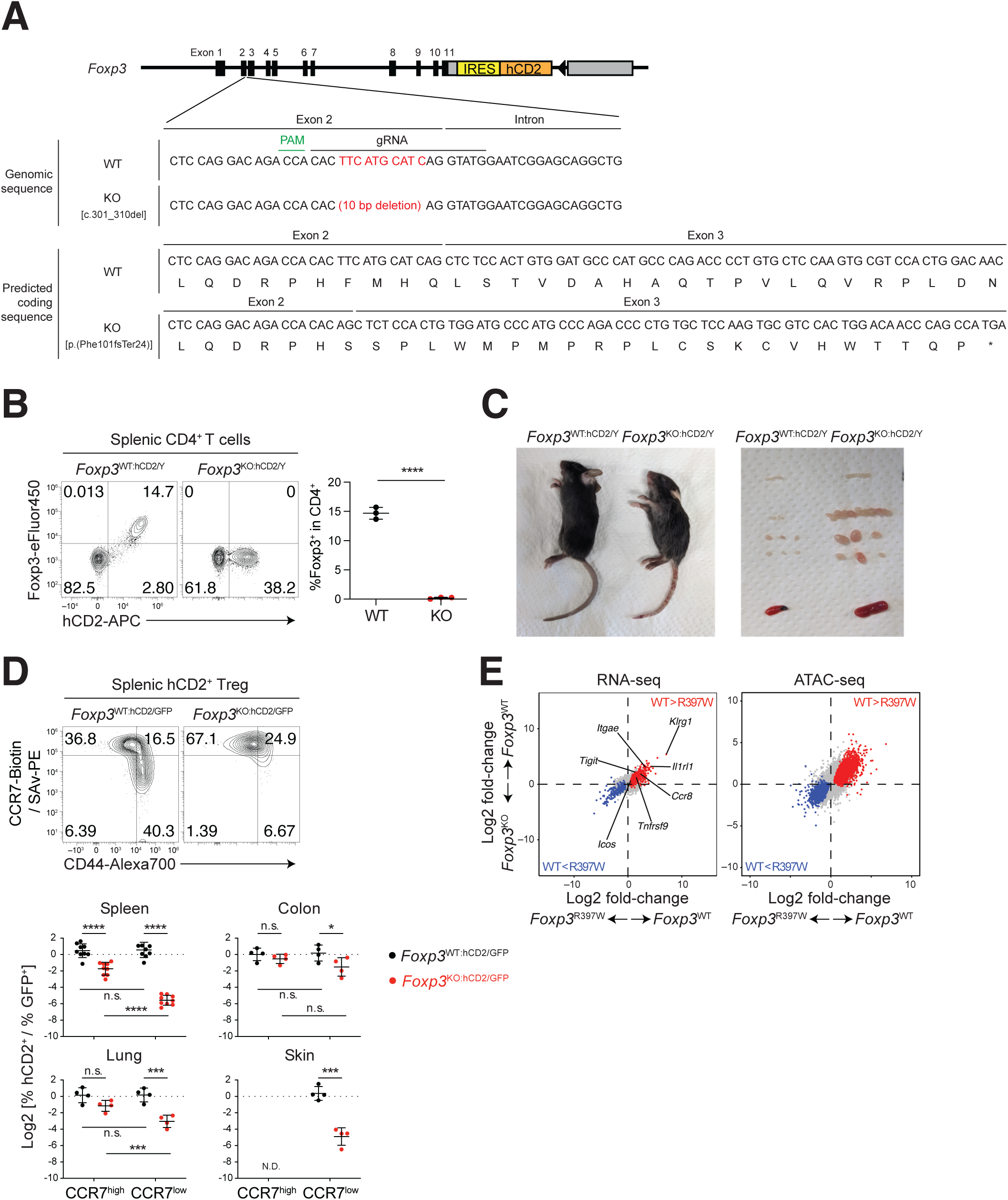
Generation and characterization of Foxp3^KO:hCD2^ mice, related to Figure 2. (A) Targeting strategy. Nucleotide and predicted amino acid sequences of the *Foxp3*^WT:hCD2^ and *Foxp3*^KO:hCD2^ alleles around the target site are shown. PAM: Protospacer adjacent motif; gRNA: guide RNA. (B, C) Analysis of three-week-old *Foxp3*^KO:hCD2/Y^ and *Foxp3*^WT:hCD2/Y^ mice. FCM of CD4^+^TCRβ^+^ cells with percentages of Foxp3^+^ cells among CD4^+^TCRβ^+^ cells summarized (*n*=3) (B). Representative images of mice, spleens, and LNs (C). (D) FCM of hCD2^+^GFP^-^ Treg cells from *Foxp3*^KO:hCD2/GFP^ or *Foxp3*^WT:hCD2/GFP^ mice. Graphs show log2 ratios of hCD2^+^ to GFP^+^ cells within CCR7^high^ or CCR7^low^ CD4^+^TCRβ^+^ populations (*n*=4–9). (E) FC vs. FC plots comparing effects of the Foxp3 R397W mutation and Foxp3 deficiency on gene expression or chromatin accessibility in Treg cells (*n*=2). Foxp3-induced (red) and Foxp3-repressed (blue) genes or OCRs (defined in Figure 2B) are highlighted. Data analyzed by unpaired t-test (B) or two-way ANOVA with Sidak’s multiple comparisons test (D).

**Figure S4:**
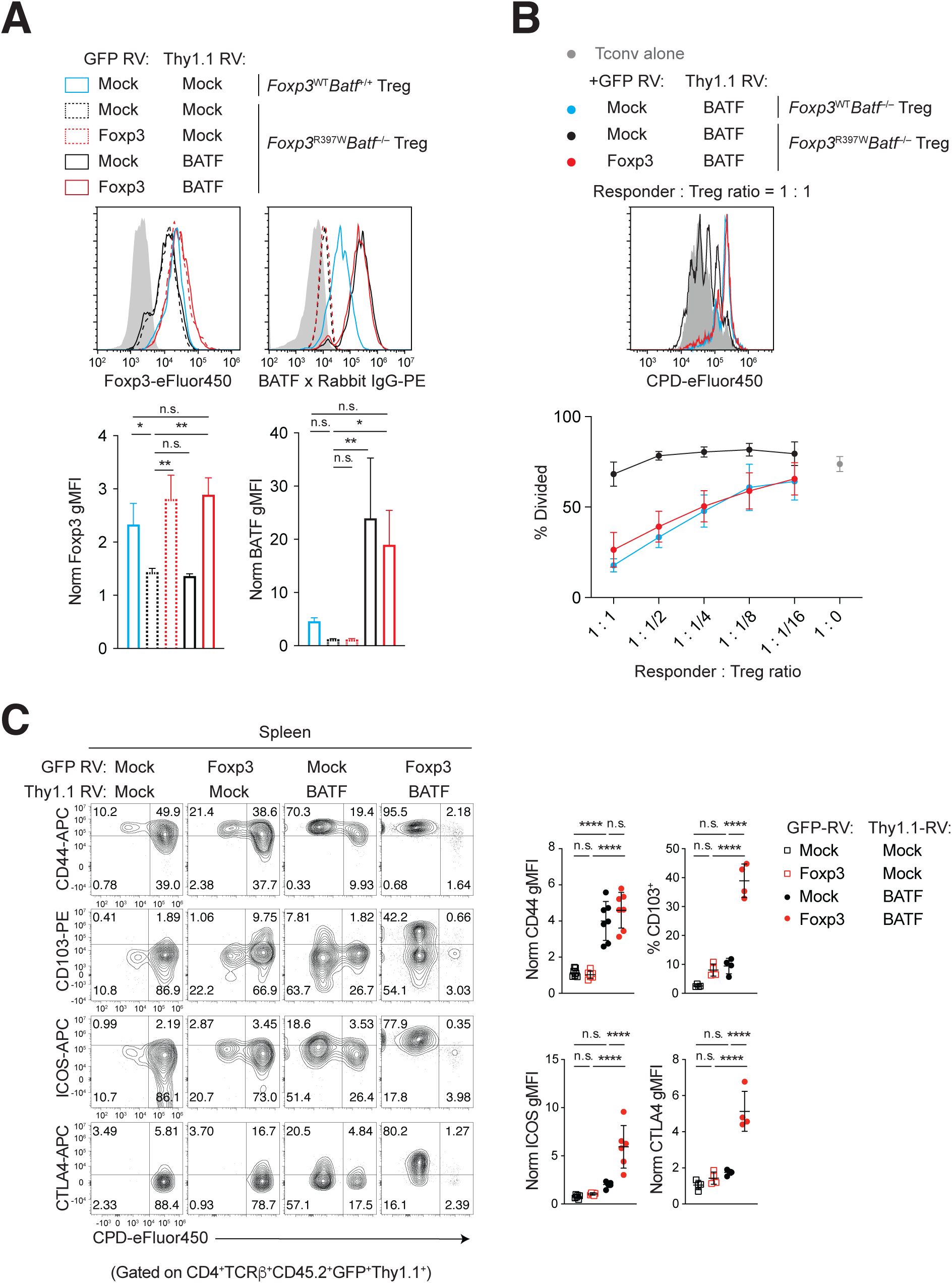
Characterization of BATF-deficient R397W Treg cells transduced with BATF and/or Foxp3, related to Figure 3. (A, B) hCD2^+^CD4^+^ Treg cells from pooled spleens and LNs of indicated mice were retrovirally transduced with the indicated RV. (A) GFP^+^ and GFP^-^ cells were sorted and analyzed by FCM. FCM of GFP^+^Thy1.1^+^hCD2^+^CD4^+^TCRβ^+^ cells, with gMFI values of Foxp3 or BATF summarized (*n*=3). gMFI values are normalized to mock RV GFP-Thy1.1-CD4^+^TCRβ^+^ cells from *Foxp3*^R397W:hCD2/Y^*Batf*^−/–^ mice in each experiment. (B) CPD eFluor 450-labeled Ly5.1 hCD2-CD4^+^ T cells were mixed with sorted Ly5.2 GFP^+^Thy1.1^+^hCD2^+^CD4^+^ cells at indicated ratios and stimulated. Representative CPD histograms of Ly5.1^+^Ly5.2-CD4^+^ T cells and percentages of divided cells are shown (*n*=3). (C) hCD2^+^CD4^+^ cells from Ly5.2 *Foxp3*^R397W:hCD2/Y^ *Batf*^−/–^ mice were retrovirally transduced, labeled with CPD eFluor 450, and transferred into irradiated Ly5.1 or Ly5.1/5.2 *Foxp3*^hCD2/Y^ mice. FCM of GFP^+^Thy1.1^+^ donor cells on day 5 post-transfer, with percentages of CD103^+^ cells and normalized gMFI of indicated molecules summarized (*n*=4–7). gMFI values are normalized to mock-transduced cells. Data analyzed by one-way ANOVA with Tukey’s multiple comparisons test (A, C).

**Figure S5:**
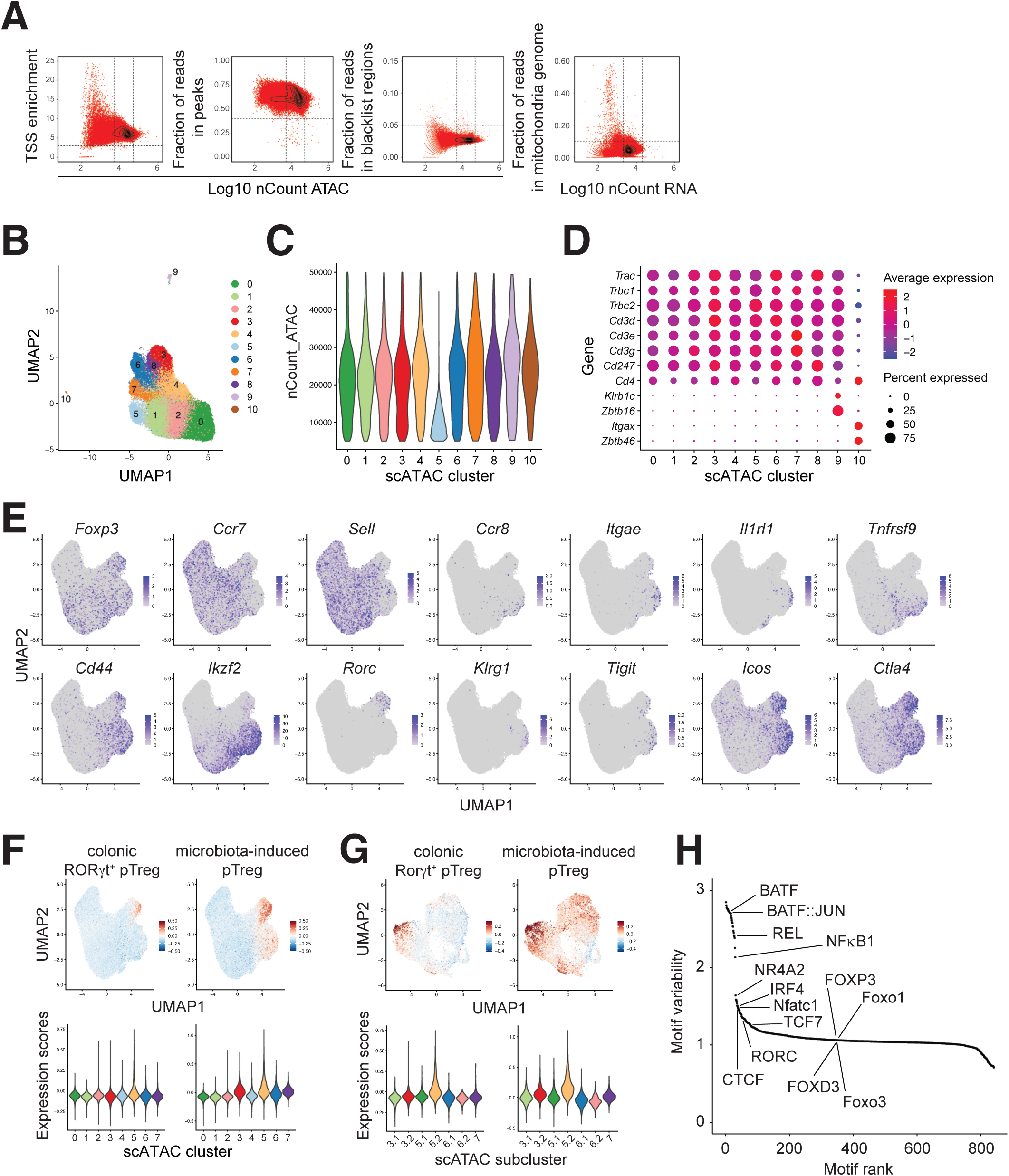
Quality control and additional analysis of scMultiome data, related to Figures 4 and 5. (A) Distribution of quality control metrics per cell. Contour plots indicate barcode density, with red dots representing cell barcodes. Dashed lines denote high-quality cell criteria. nCount ATAC: unique molecular identifiers (UMIs) per cell barcode for scATAC-seq libraries; nCount RNA: UMIs per cell barcode for scRNA-seq libraries; TSS: transcription start site. (B) Merged UMAP of scATAC-seq data after filtering and data integration with batch effect correction, prior to outlier cluster removal. (C) Violin plot of nCount ATAC per cluster. Cluster 5, with low nCount ATAC, was excluded from further analysis. (D) Dot plot of cluster-level expression of indicated genes. Cluster 9 represents NKT cells based on *Klrb1c* (NK1.1), *Zbtb16* (PLZF), and αβTCR complex genes, and cluster 10 represents conventional dendritic cells based on *Itgax* (CD11c) and *Zbtb46*, lacking αβTCR complex genes. These clusters were excluded from further analysis. (E) Single-cell-level expression of indicated genes. (F, G) Gene expression scores for gene signatures of colonic RORγt^+^ Treg cells^36^ and microbiota-induced pTreg cells^37^, visualized in UMAPs for all cells (F) or eTreg and Teff cell subclusters (G). Violin plots show scores per cluster (F) or subcluster (G). (H) Variability of TF motifs across individual cells, ranked by variability, with key TFs highlighted.

**Figure S6:**
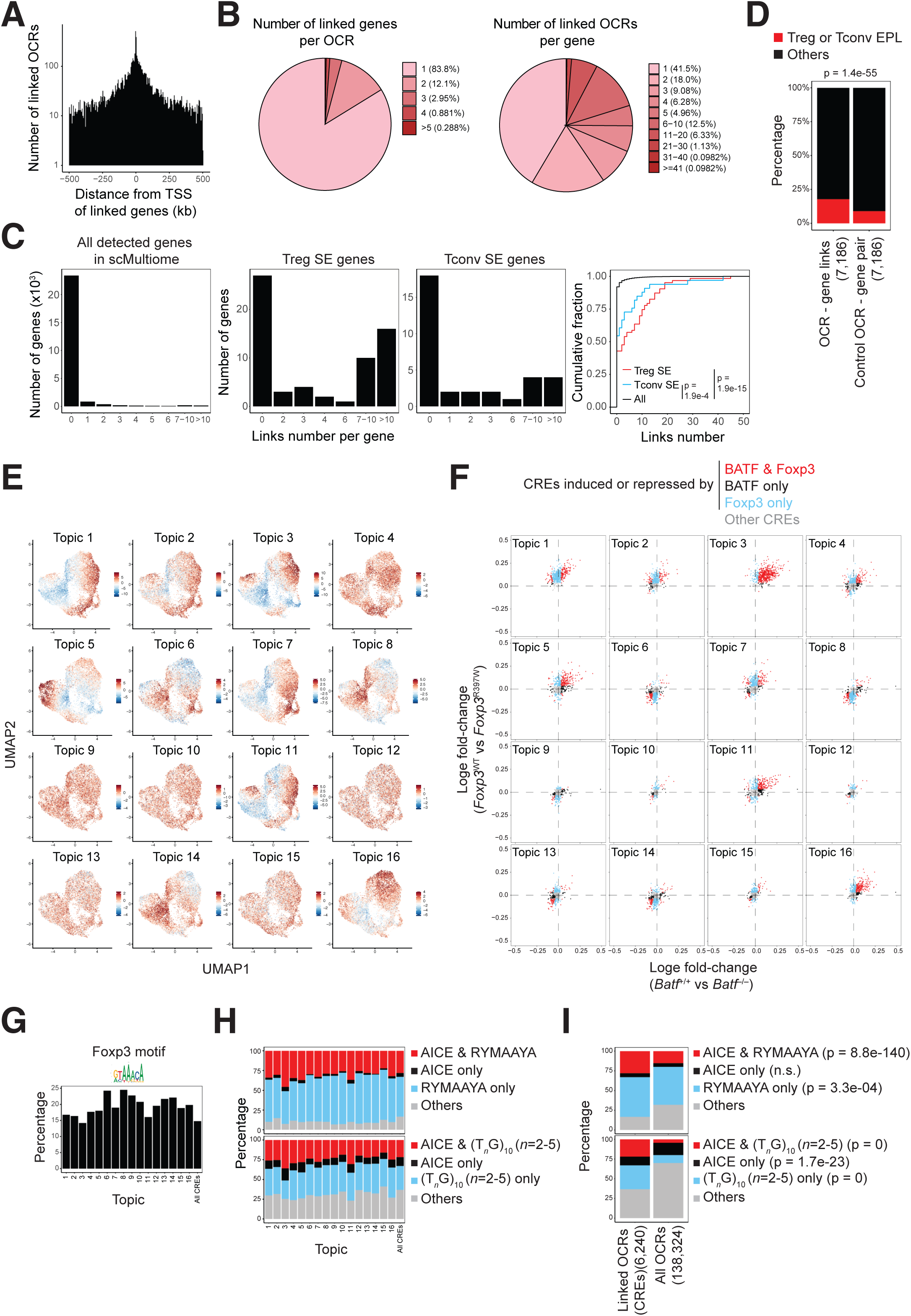
Characterization of OCR-gene links and cis-regulatory topics, related to Figure 6. (A) Distribution of distances between OCRs and TSSs for OCR-gene links. (B) Frequencies of numbers of genes linked to each OCR or OCRs linked to each gene. (C) Distribution of OCR-gene links per gene for all detected genes, Treg SE genes, and Tconv SE genes, as defined by H3K27ac ChIP-seq^41^. CDF plot shows the cumulative distribution of OCR-gene links for these gene categories. (D) Percentages of OCR-gene links or control OCR-gene pairs overlapping with enhancer-promoter loops (EPLs) detected in Treg and/or Tconv cells, as defined by H3K27ac HiChIP^42^. Control pairs are randomly selected and matched for OCRs-TSS distances. (E) Accessibility scores for topic-associated CREs in eTreg and Teff cell subclusters, visualized on UMAPs. (F) FC vs. FC plots comparing the effect of BATF deficiency and the R397W mutation on chromatin accessibility of topic-associated CREs. Each dot represents a CRE. (G, H) Percentages of CREs with the canonical Foxp3 motif (MA0850.1) (G) or indicated combinations of TF motifs (H) among topic-associated or all CREs. (I) Percentages of OCRs with indicated combinations of TF motifs in CREs and all OCRs. Data analyzed by Kolmogorov-Smirnov test (C) or Fisher’s exact test (D, I).

**Figure S7:**
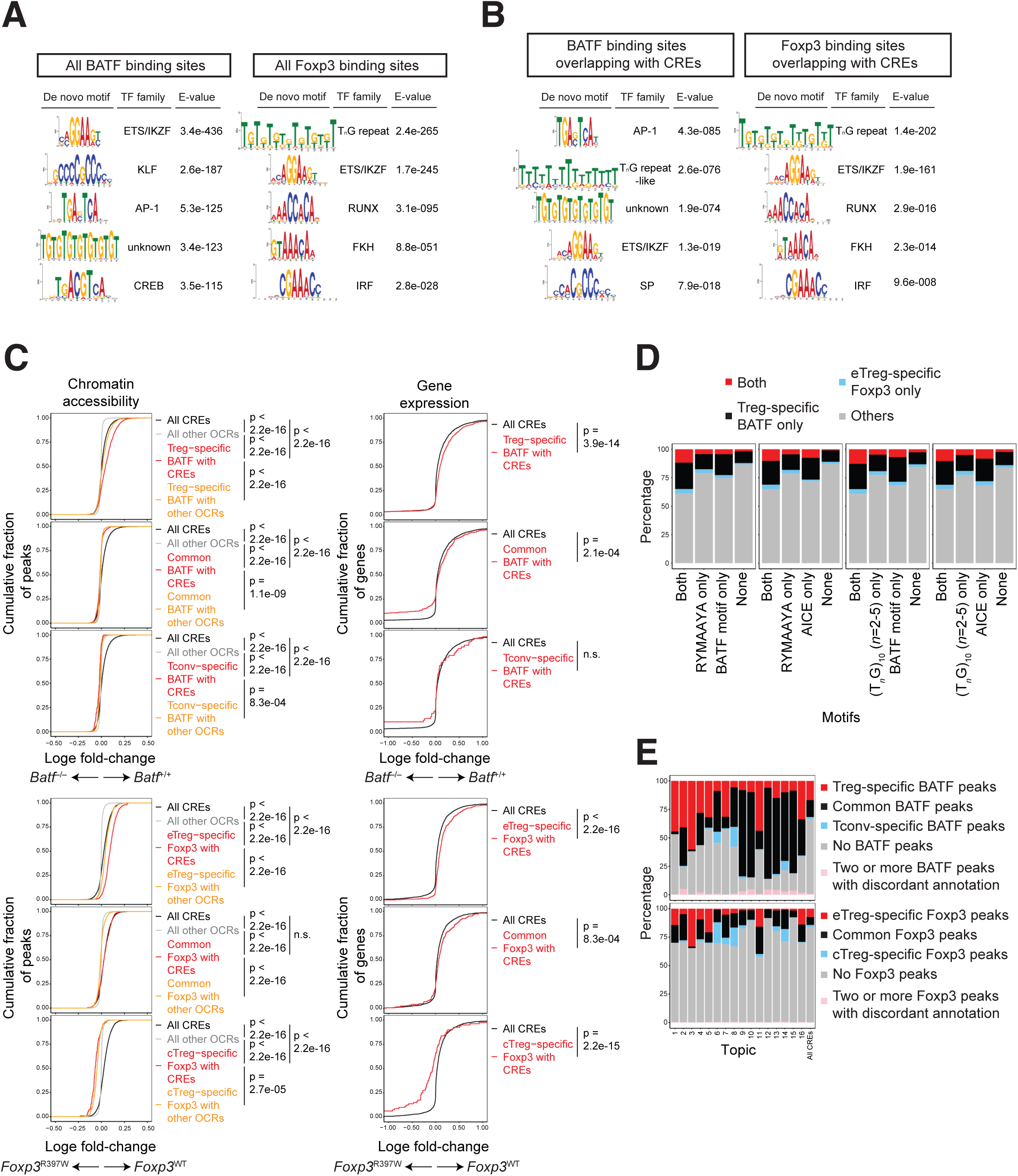
Characterization of BATF and Foxp3 binding sites, related to Figure 7. (A, B) Top five *de novo* motifs significantly enriched in all BATF or Foxp3 binding sites (A) or in BATF or Foxp3 binding sites overlapping with CREs (B). (C) CDF plots showing effects of BATF deficiency or the Foxp3 R397W mutation on accessibility of CREs or other OCRs overlapping with indicated categories of BATF or Foxp3 binding sites, and on expression of associated genes. (D) Percentages of CREs overlapping with indicated categories of BATF and/or Foxp3 binding sites in CREs with specified TF motif combinations. (E) Percentages of CREs overlapping with indicated categories of BATF and/or Foxp3 binding sites in topic-associated or all CREs. Data analyzed by Kolmogorov-Smirnov test (C).

## Key resources table

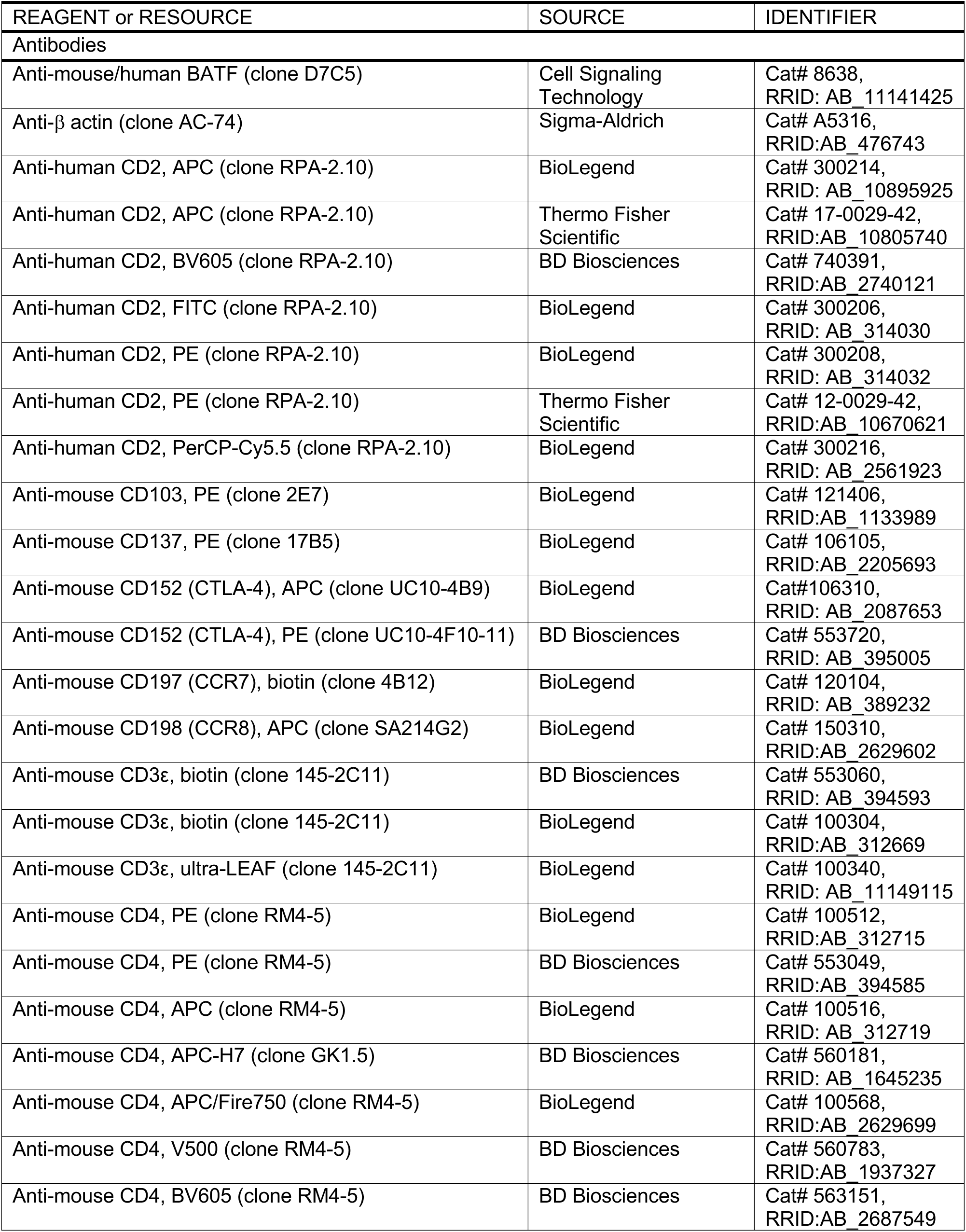
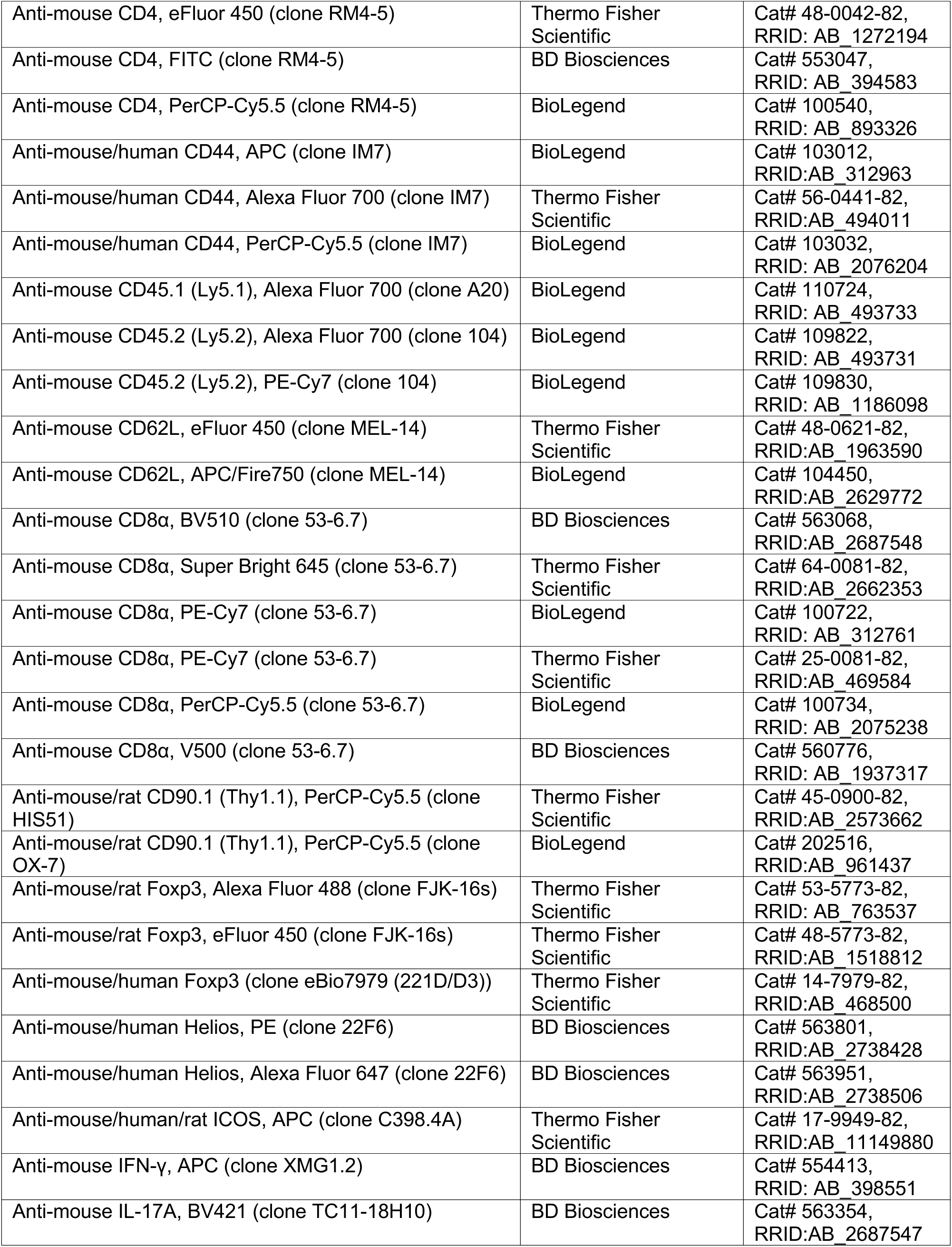
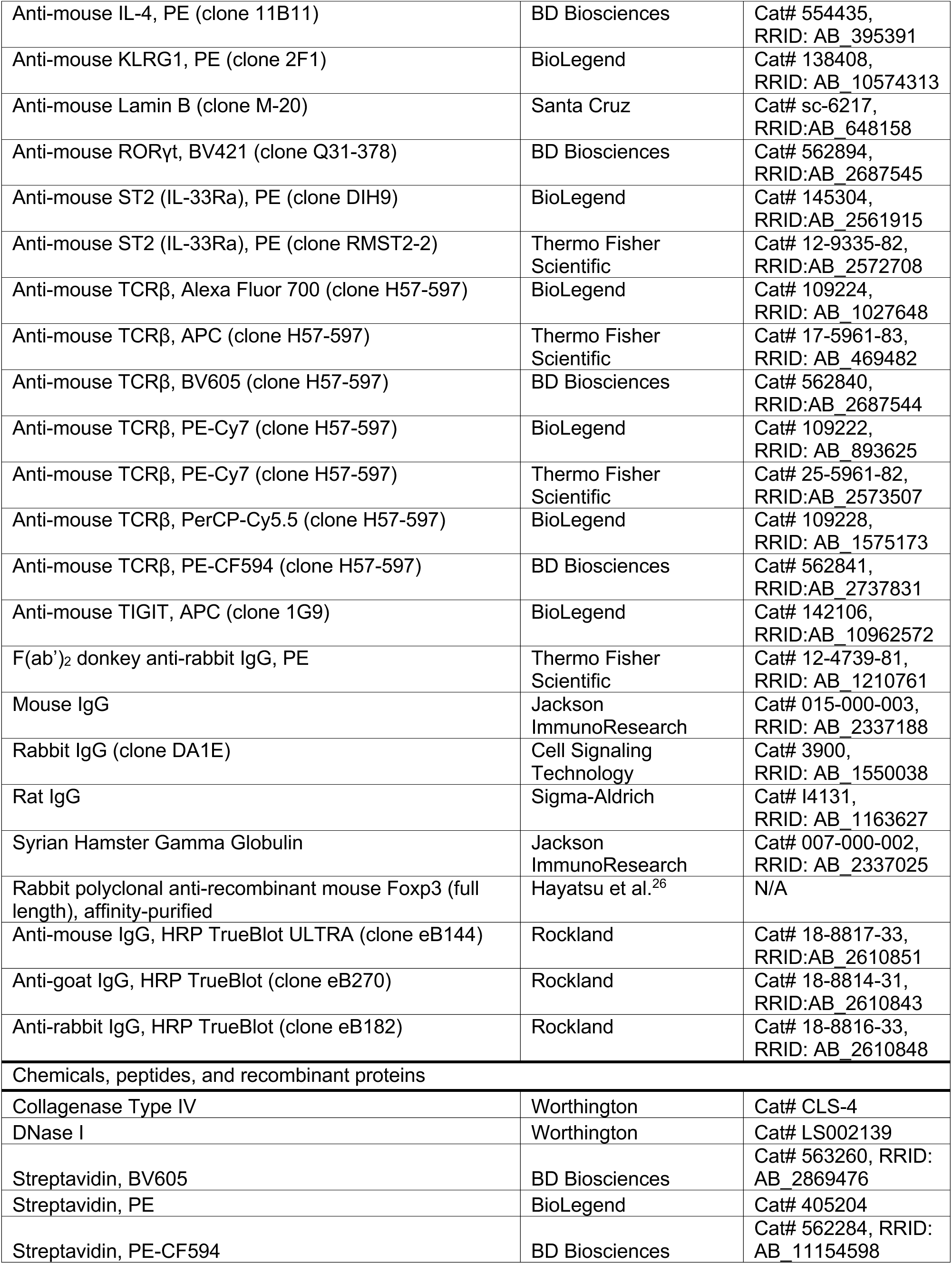
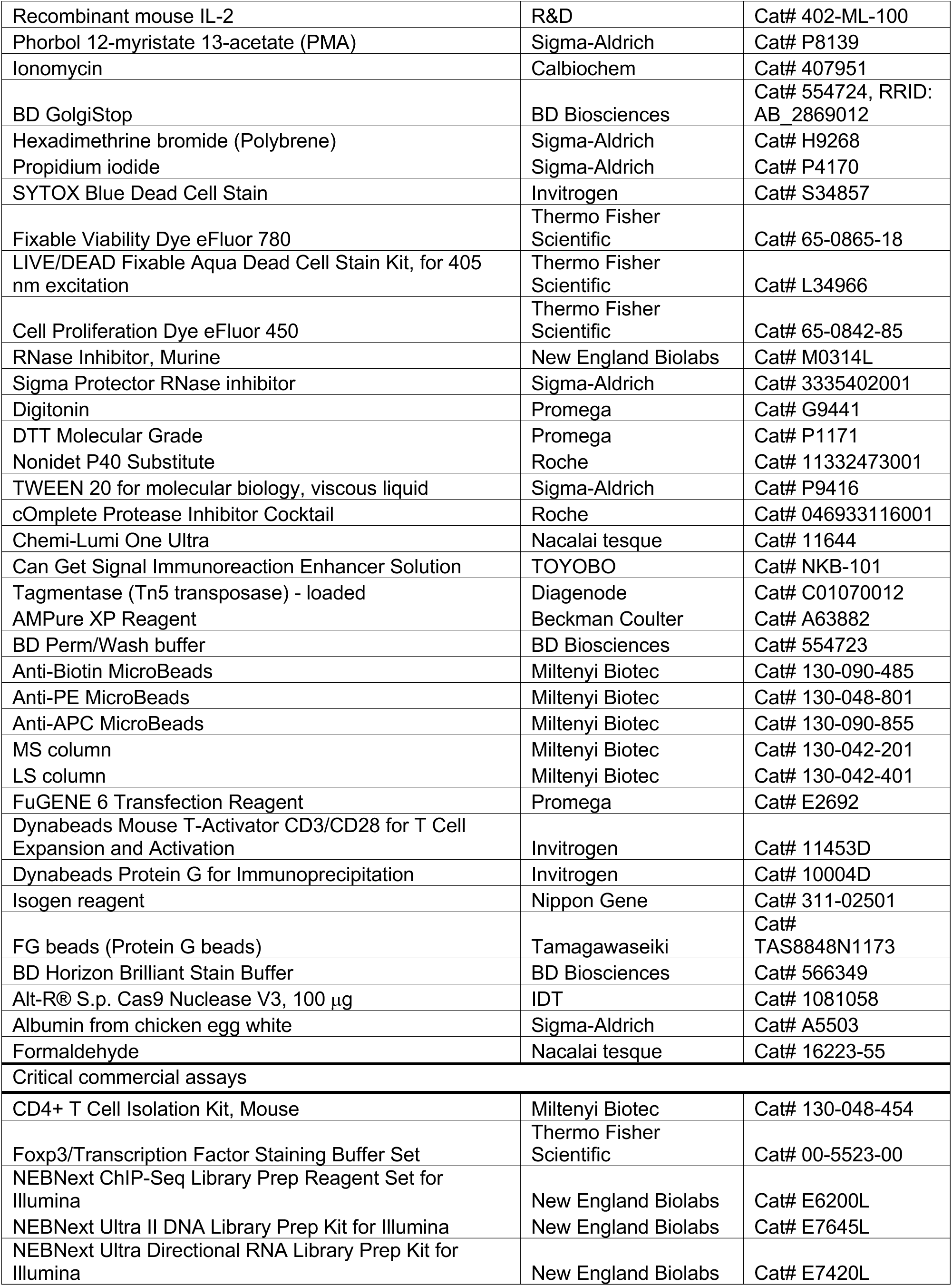
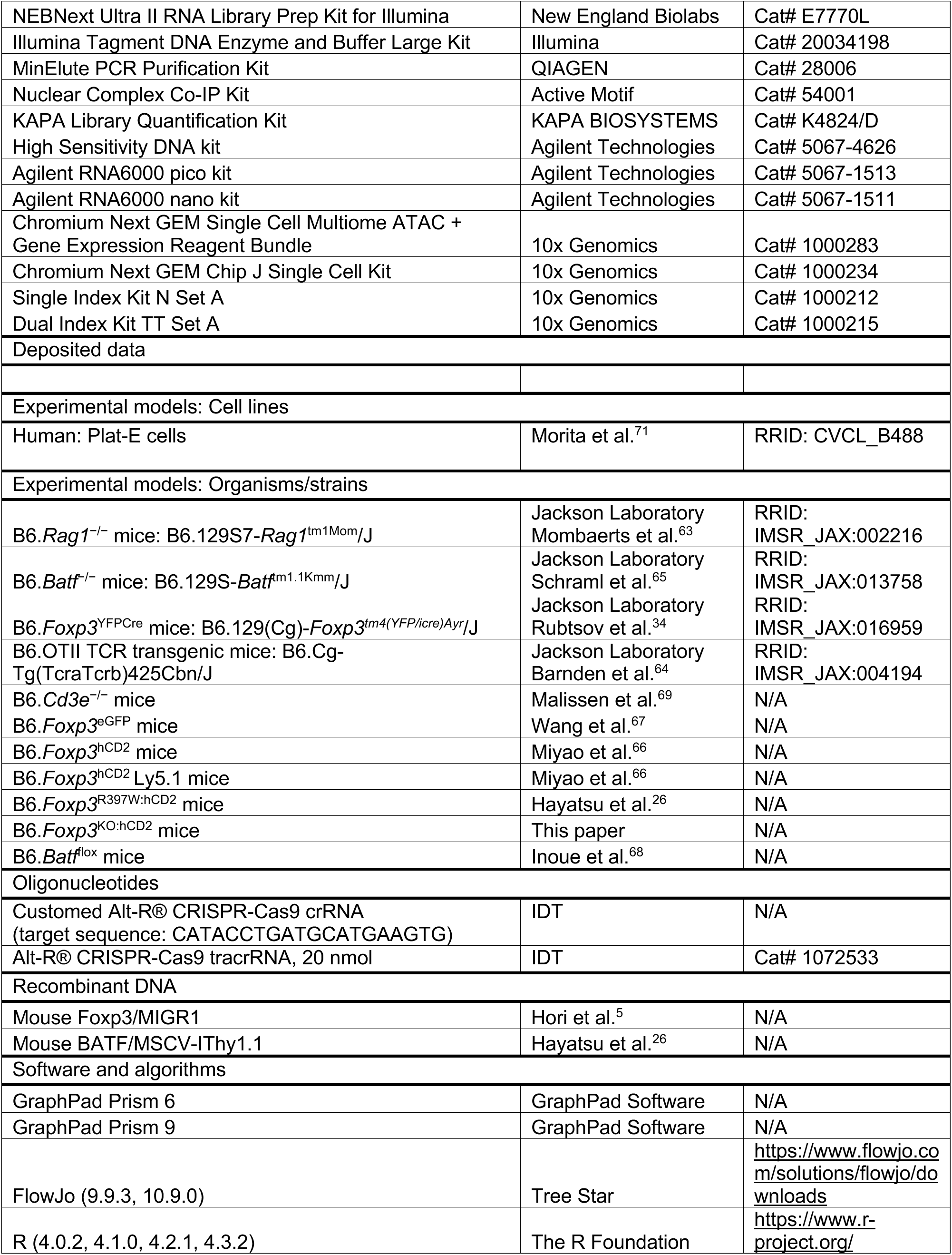
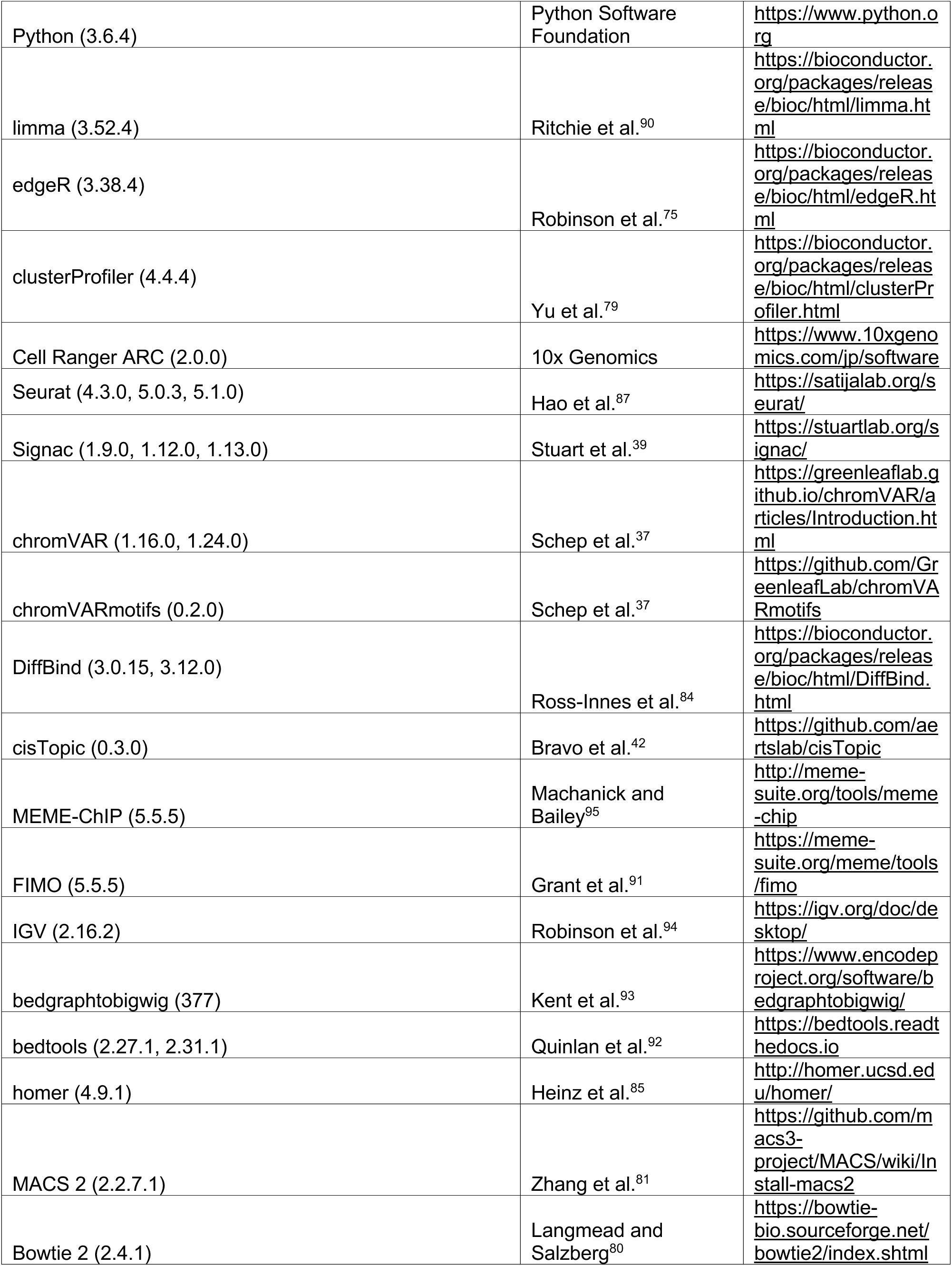
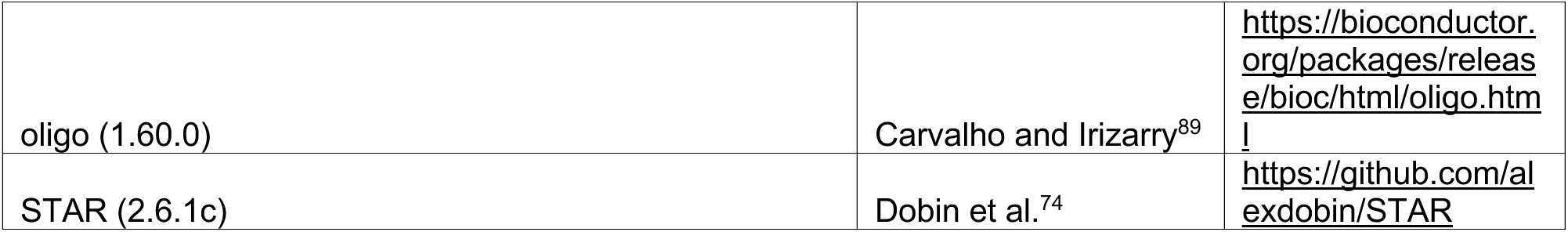

